# Exploring individual codon influence on protein expression: a predictive approach

**DOI:** 10.1101/2024.06.30.601029

**Authors:** Konstantin Zaytsev, Natalya Bogatyreva, Alexey Fedorov

## Abstract

An important role of a particular synonymous codon composition of a gene on its expression level is well-known. There are a number of algorithms optimising codon usage of recombinant genes to maximise their expression in host cells. Nevertheless, the problem has not been solved yet and remains relevant. In the realm of modern biotechnology, directing protein production to a specific level is crucial for metabolic engineering, genome rewriting, and a growing number of other applications. In this study, we propose two new simple statistical and empirical methods for predicting the protein expression level from the nucleotide sequence of the corresponding gene: Codon Expression Index Score (CEIS) and Codon Productivity Score (CPS). Both of these methods are based on the influence of each individual codon in the gene on the overall expression level of the encoded protein and the frequencies of isoacceptors in the species. Our predictions achieve a correlation with experimentally measured quantitative proteome data of *Escherichia coli* up to a level of r=0.7, which is superior to any previously proposed methods. Our work helps to understand how codons determine translation rates; based on our methods, it is possible to design proteins optimised for expression in a particular organism.

## INTRODUCTION

Most amino acids are represented by two or more codons, and mutations that substitute one codon for another synonymous codon do not alter the amino acid in the gene’s final product. Traditionally viewed as silent, these mutations were believed to have minimal impact on phenotype. However, recent research has uncovered various instances where synonymous mutations play significant roles (1–3). These roles include optimising gene expression by boosting translation initiation, adjusting translation speed by influencing codon usage and mRNA structures, stabilising mRNA to prevent premature degradation pre-translation, and affecting protein folding, degradation, ubiquitination, and protein secretion within cells (3). One of the most impressive experiments showed the effect of synonymous mutations in GFP on a 250-fold change in expression levels (4). Experiments with TEM-1 β-lactamase suggest that synonymous mutations may have beneficial effects by increasing the expression of an enzyme with low substrate activity (5). In experiments on *Salmonella enterica*, it was shown that the effects of synonymous mutations are due to a combination of effects on mRNA stability and translation efficiency, which alter levels of the weak-link enzyme. These studies indicate that synonymous mutations most likely play an underestimated role (6). Understanding the mechanism of synonymous mutation impact is important for understanding of evolution and variation.

Multiple studies showed that the amount of expressed protein could be increased by replacing codons in a gene with the synonymous ones (7–11). However, replacing all codons with the preferred ones does not necessarily achieve the maximum protein yield. Instead, it can reduce the balance between codon usage and tRNA abundance, which leads to reduced global translation efficiency (12). Other possible consequences include changes in protein solubility (13) and incorrect folding (14–16).

The ability to change a gene sequence to modify protein expression to the predetermined level is an important industrial and academic challenge. Metabolic engineering often requires insertion of groups of genes to be expressed with specific individual levels to provide maximal efficiency of the required set of enzymatic reactions while avoiding unnecessary energy cost for production of excessive amounts of proteins (17). Another trending issue is genome rewriting (18–20), which requires changes in synonymous codon sets of protein-encoding genes and may lead to imbalanced production of proteins that could harm cell metabolism.

The most popular methods for assessing expression levels are CAI (Codon Adaptation Index) (21) and TAI (tRNA adaptation index) (22–24). TAI predicts the protein expression based on the pool of available tRNAs and the binding efficiency for codons with corresponding isoacceptor tRNAs. Its expression prediction accuracy was tested on E. coli genes at r=0.54 level (24). However, this method does not take the mRNA stability into account, which does also depend on codon usage (25). The Codon Adaptation Index (CAI) (21) is the most widely used method for predicting the gene expression level from its sequence. The method is based on determining organism’s preferred codons for each amino acid from their occurrence in well-expressed genes. CAI for a gene is calculated as the geometric mean of the frequency of each codon occurrence relative to the most frequent synonymous codon in the training set. The main advantage of this method is that it is very efficient in terms of requiring a fairly small amount of data for training: only a few sequences of well-expressed genes. However, CAI is based on the assumption that there is only one preferred codon for each amino acid, and the higher the proportion of the preferred codons, the higher the level of protein expression is. CAI expression prediction accuracy was tested on 96 *E. coli* genes at r=0.53 level (9). Another method for expression prediction is the Relative Codon Bias Score (RCBS) (10). It measures codon frequency bias by comparing codon frequencies to the frequencies of the individual nucleotides. Authors claimed r=0.7 expression prediction accuracy, after testing on a dataset, consisting of 45 *E. coli* genes. There are also numerous other indices and methods, which uncover links between codon usage and different aspects of gene expression (26). Even though these indices were trained on different data sources, they tend to correlate with each other.

Due to advancements in biotechnology, there are several large scale proteome analysis datasets for different organisms (27–33). This allows us to create more powerful methods for expression prediction based on genome-wide analysis and expand the understanding of expression regulation patterns. The goal of this study has been to show the influence each of the individual codons has on the integral level of protein expression. This would allow better optimisation for expression of recombinant genes, as well as their alteration to the desired expression level.

## MATERIAL AND METHODS

### Dataset

In this study, we analysed levels of protein expression for *E. coli* genes. By protein expression we mean the number of protein copies produced from the corresponding gene, which is a function of both transcription and translation. Genomic sequence for *E. coli* strand ATCC 25922 and all the open reading frames (list of genes) were obtained from [https://genomes.atcc.org/genomes/ccbc9e61ad334c2c].

In this study we used protein abundance data for *E. coli* from (27) where peptides were analysed by LC-MS/MS in three repetitions. Protein content per bacterial cell was calculated on the basis of the DNA content. For protein expression values, we used the average of the 3 experimental values. The range of non-zero expression values in the dataset varied from 0.06 to 38022 protein copies per cell. The intersection of expression data and multiple nucleotide sequences of genes was performed using the UniprotID of the encoded proteins. Only genes with exactly 1 copy in the genome and non-zero expression levels were selected for further analysis. In total, the dataset we used included 1688 genes.

### Gene clustering

Based on the expression level, we divided genes into four classes: genes with the highest expression level were combined into Class 1, genes with the lowest one were combined into Class 4. Genes with the higher and lower than median expression levels were combined into Class 2 and Class 3 respectively. Each of the obtained classes contained 422 genes.

For each of four classes we calculated frequencies of codon appearances for all amino acid coding codons. Because gene lengths were different and so was the total number of codons in each class, we normalised codon distributions for the classes, so that they would sum up to 1.

To show the difference between *E. coli* genes and genes from other organisms, we created a class, consisting of non-*E. coli* genes. We used a dataset with genes originating from different organisms, that were expressed in *E. coli* (34). From the group of such genes, that were producing the highest amount of protein, we randomly selected 422 genes and combined them into the Alien Class. Frequencies of codon appearances in the class were also normalised so that they sum up to 1.

### Codon expression index (CEI)

We used Kendall’s rank correlation to determine the influence of each of the codons on the protein expression level (35). The essence of rank correlation lies in the direction of changes between the values, rather than the specific ratios of values.

For each codon *c*, except for the stop codons, we created an array of its frequencies in the genes from the dataset 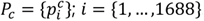. This way we got 61 arrays (number of amino acid coding codons) with 1688 values (number of genes in the dataset) in each one. We also created an array with numbers of protein copies per cell produced by genes from the dataset *E* = {*e_i_*}. This array also contained 1688 values. Array indices were unified for all the arrays and represented specific genes. We calculated Kendall’s tau between the frequency array for each of the codons and the array of expression values.

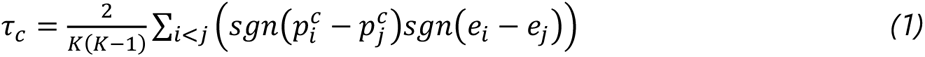

Here *K* is the number of genes in the dataset. *τ_c_* is the coefficient of Kendall’s rank correlation. For each correlation coefficient, we estimated its statistical significance *Z* (36).

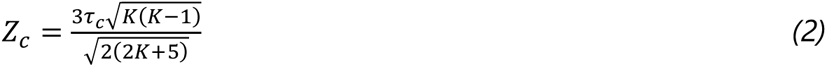

*Z_c_* shows the number of standard deviations by which the resulting value differs from what would be expected if both arrays were totally independent. Absolute *Z* values greater than 3 are considered significant. This enabled us to determine the presence of a positive or negative influence of the frequency of a given codon on the overall expression. Additionally, the sign of *Z* values indicates a positive or negative correlation. To test this, we calculated a set of *Z* values for correlations between codon frequency array and a randomly shuffled expression array. All obtained *Z* values fell within the range from −3 to 3, confirming the selected statistical significance threshold. We named the set of *Z* values for codons as Codon Expression Index (CEI).

### Codon expression index score (CEIS)

We used the obtained CEI values for creating a protein expression prediction model. We sorted genes from the dataset in the decreasing order of expression and split the dataset into two disjoint sets: training and test. Every 10 out of 11 genes were selected into the training set and every 11^th^ gene was selected into the test set.

CEI values {*Z_c_*}; *c* = {1,… ,61} were calculated from the training set. Then for each gene from the training set, we calculated the average CEI value of its codons and combined the results into an array 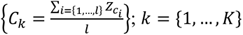, where *K* is the number of genes in the training set; *l* is the number of codons in a gene. We created a second array containing logarithmic protein expression values. To eliminate negative log values for the genes with less than 1 protein copy per cell, we added 1 to each of the expression values before taking the logarithm: *log E* = {*log*(*e_k_* + 1)}. From these two arrays, we calculated linear regression coefficients *a* and *b* using least squares approximation. These coefficients allowed us to scale the predicted values to the actual expression level: *P_k_* = *aC_k_* + *b*.

The model was then tested on the test set. To better test the prediction accuracy, we repeated the dataset split into training and test sets 10 more times, so that each gene from the dataset appeared in the test set exactly once. For each pair of sets, we calculated CEI values and regression coefficients from the training set, and then predicted expression values *P_k_* for genes from the test set. All the predicted values were combined into an array. Actual expression values *E_k_* were combined into second array. We used Pearson’s coefficient of linear correlation between the arrays of predicted and actual expression values as a measure of prediction accuracy.

It is important to note that the use of regression did not affect the prediction accuracy, but it improved the convenience of interpretation of the predicted expression values.

### Codon productivity (CP)

For each codon *c*, we calculated the total number of the corresponding amino acids produced by the cell.

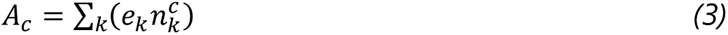

Here, *e_k_* is the number of protein copies per cell produced from the gene *k*. 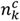 is the number of occurrences of codon *c* in the gene *k*.

Next for each codon *c* we calculated its total number of appearances in the genes from the dataset.

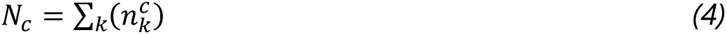

Codon productivity (CP) was calculated by dividing the number of amino acid copies produced by each codon by the number of codon appearances.

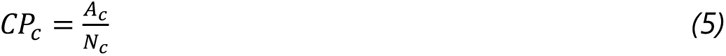

CP is the average number of amino acids produced by the cell from a single codon.

### Codon productivity score (CPS)

The model for expression prediction based on CP values is similar to the one for CEI values, except for the use of the average CP for the gene instead of the average CEI.

### Codon adaptation index (CAI)

For Codon adaptation index (CAI) (21) calculations, we used the CAI v.1.0.5 module for Python (37) [https://github.com/Benjamin-Lee/CodonAdaptationIndex]. Prediction accuracy was tested using 11-fold cross-validation, same method as with CEIS and CPEIS. As a reference set of genes, required by CAI we used a quarter of genes from the training set with the highest expression.

### tRNA adaptation index (TAI)

For tRNA adaptation index (TAI) (22) calculations, we used the codon-bias v.0.3.1 module for Python [https://github.com/alondmnt/codon-bias], [https://doi.org/10.5281/zenodo.8039452]. The tRNA gene copy numbers we used, were acquired from [http://gtrnadb.ucsc.edu/GtRNAdb2/genomes/bacteria/Esch_coli_ATCC_25922/]. This method did not require any reference sequences, so we calculated TAI values for all the genes from the dataset and compared them with actual expression values using Pearson’s linear correlation.

### Relative codon bias score (RCBS)

For the Relative codon bias score (RCBS) (10) calculations, we also used the codon-bias v.0.3.1 module for Python [https://github.com/alondmnt/codon-bias], [https://doi.org/10.5281/zenodo.8039452]. All parameters were set to default values. This method also did not require any reference sequences, so we calculated prediction accuracy the same way as for TAI.

## RESULTS

The structure of each gene is unique and depends on the structure and functions of the protein it encodes. To show the differences in the codon structures of genes coding for highly- and low-expressed proteins, we combined genes from the dataset into four classes based on the levels of the corresponding protein production. The first class contained genes with the highest expression, and the fourth contained genes with the lowest one. We calculated distributions of codon frequencies for the classes, which allowed us to minimise the individual influence of each gene on the codon distribution while maintaining factors that influence the overall expression level.

Genes were split into the classes in such a way that each class contained the same number of genes; therefore, boundaries of the classes have no biological meaning. Expression levels of the genes from the Class 1 ranged from 356 to 38022 protein copies per cell, Class 2: from 82 to 356, Class 3: from 18 to 82, Class 4: from 0 to 18.

Figure 1 shows that for some of the codons, their frequency of occurrence does not depend on the levels of protein expression (for example, AGG, TGT, CCT, CCA, GAG, ATG, CAG, etc.), while for other codons, the differences are significant (for example, TCT, GCT, TTA, GTT, GGT, TTT, AAA, etc.). We calculated frequencies of codon occurrences as fractions of all 61 possible codons occurrences in the genes from the particular class. As can be seen, all differences between codon frequencies are directional and proportional to the differences between average expression levels of the classes. Therefore, it is obvious that there is a link between the frequency of each individual codon and the integral protein expression level.

**Figure 1.**
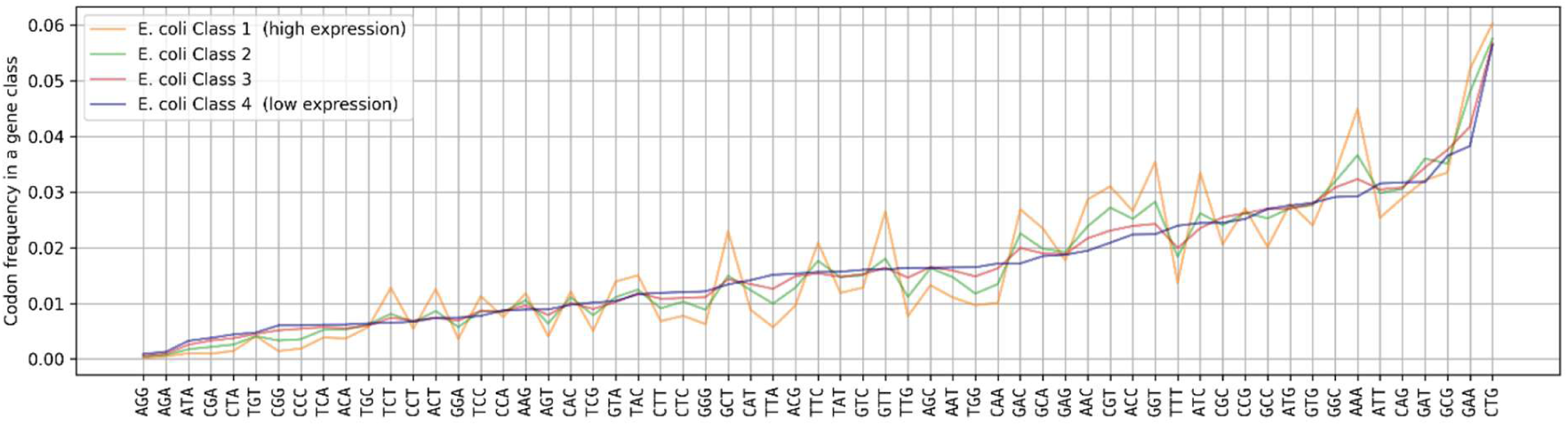
The distribution of codon frequencies in 4 classes of *E. coli* genes with different levels of protein expression. Codons are sorted by their frequency of occurrence in the class of genes with the lowest expression. The class of genes with the highest expression levels is shown in orange (Class 1), the class of genes with the lowest expression levels is shown in blue (Class 4), classes of genes with intermediate expression levels are shown in green (Class 2) and red (Class 3).

Figure 2 shows distributions of codon occurrences in individual genes from Class 1 (highest expression genes) and Class 4 (lowest expression). Distributions depend on the number of genes containing such codon and their frequency of appearances in genes. *E. coli* exhibits preferences for some specific codons for several amino acids (Leucine, Isoleucine, Arginine, Glycine), or synonymous codons can be used equally (for example for Phenylalanine). But there is always difference in codon preferences between the genes with different levels of protein expression. What remains consistent across all codons is that the codon occurrence distribution for the genes with low expression resembles the normal distribution much better than for the genes with high expression, where there is much higher specificity of codon frequencies.

**Figure 2.**
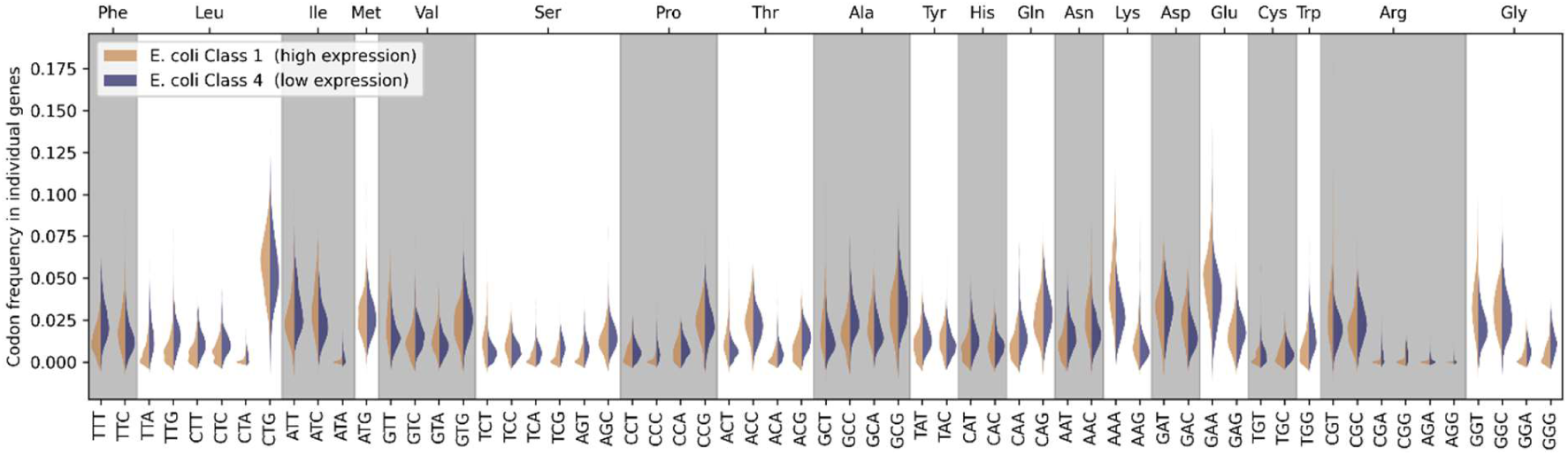
Distributions of codon appearance frequencies in the individual *E. coli* genes from the class of genes with the highest expression levels (Class 1) are shown in orange. Distributions for the class of genes with the lowest expression levels (Class 4) are shown in blue.

Figure 3 shows the distribution of codon frequencies in each of the four classes of *E. coli* genes and a class of alien genes, which were derived from other species. We sourced expression data for genes from different species from (34). Among 1973 alien genes with the highest expression score in the dataset, we randomly chose 422 and combined them into a single “Alien Class”. We can see that the frequencies of some codons are specific to the *E. coli* genes, while the frequencies of other codons do not show the species specificity. This leads us to assume that the codon frequency distribution is specific to a particular species, and when used to predict the level of protein expression, it makes sense to take the species-specific nature of the codon distribution into account.

**Figure 3.**
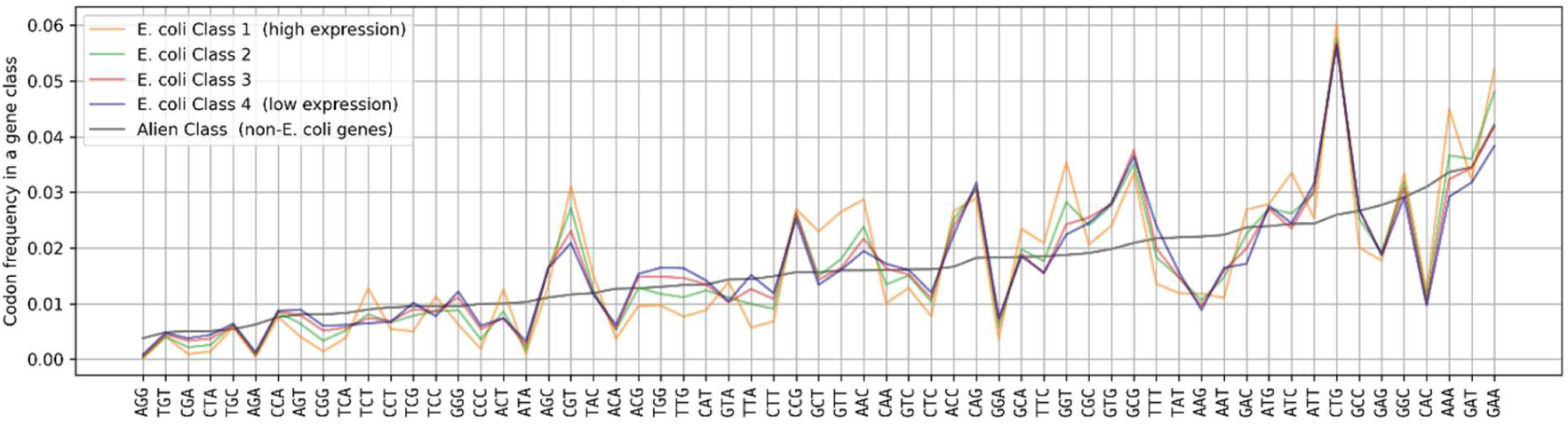
The distribution of codon frequencies in each class of *E. coli* genes and a group of genes from other species (Class of non-*E. coli* genes). Codons are sorted by the frequency of their appearance in non-*E. coli* genes, shown in grey. The class of genes with the highest expression levels is shown in orange (Class 1), the class of genes with the lowest expression levels is shown in blue (Class 4), classes of genes with intermediate expression levels are shown in green (Class 2) and red (Class 3).

Despite the fact that alien genes were expressed well in *E. coli* in many instances, their codon distribution is quite different from the distributions for native genes. We have listed linear correlation values between distributions for each pair of classes in the Table 1. The lowest observed correlation coefficient between a pair of distributions for *E. coli* gene classes is 0.89, while the correlation between *E. coli* classes and the class of foreign genes ranges from 0.73 to 0.77. So, we can talk about the presence of gene optimisation to the specific species in genes with all levels of expression. The absence of such optimisation does not necessarily impair gene expression. This also means that the effect of codon usage on the protein expression level is individual for each species.

**Table 1.** Coefficients of linear correlation between pairs of codon distributions for four classes of *E. coli* genes with different levels of protein expression and a class of non-*E. coli* genes.

|  | Class 1 (high expression) | Class 2 | Class 3 | Class 4 (low expression) |
| --- | --- | --- | --- | --- |
| Class 1 (high expression) | 1.00 | 0.96 | 0.92 | 0.89 |
| Class 2 | 0.96 | 1.00 | 0.98 | 0.97 |
| Class 3 | 0.92 | 0.98 | 1.00 | 0.99 |
| Class 4 (low expression) | 0.89 | 0.97 | 0.99 | 1.00 |
| Class of non-*E. coli* genes | 0.73 | 0.77 | 0.75 | 0.73 |

To analyse the contribution of the individual codons to the integral protein expression level, we introduced the Codon Expression Index (CEI), which shows the level of statistical significance of the correlation between the frequency of each codon occurrence in E. coli genes and the expression level of the corresponding protein. Figure 4 shows CEI values for each of the codons. Orange dots represent random CEI values, which we used to determine the statistical significance boundary. All random (orange dots) values fall within the range from - 3 to 3 (red lines), corresponding to three standard deviations. Therefore, we can assume that codons for which the CEI module is greater than 3 have a significant effect on the integral level of expression.

**Figure 4.**
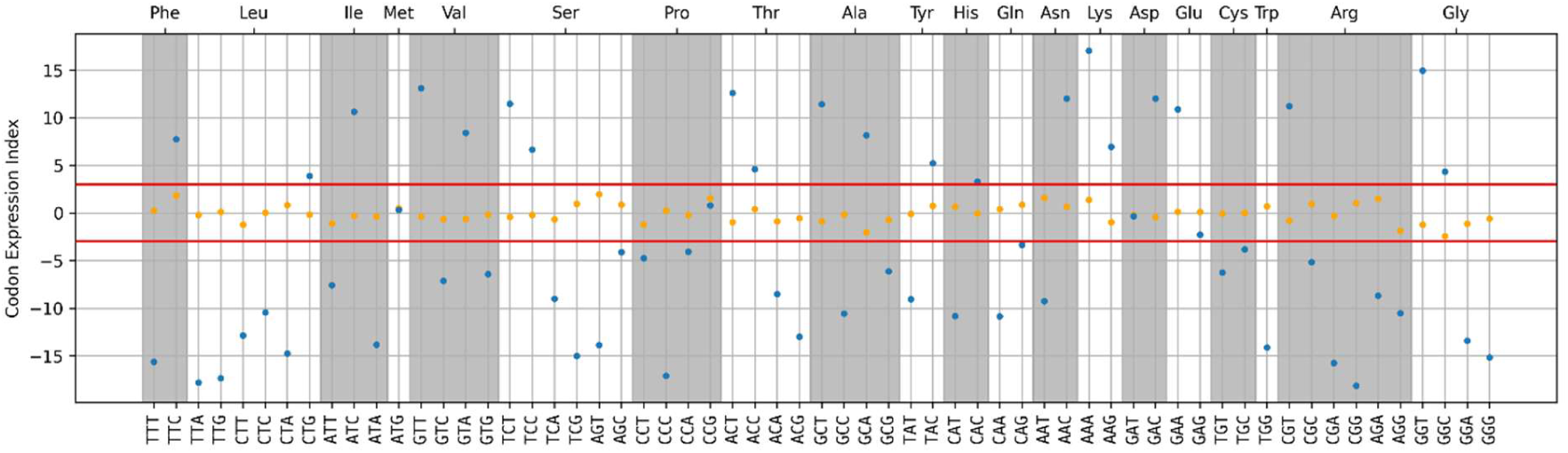
Codon expression index (CEI) values show the level of influence of a single codon on the protein expression level of the gene (blue dots). Orange dots show the CEI values for randomly shuffled arrays and are used to determine boundaries of statistical significance. All random values fall within the range from −3 to 3 (red lines), corresponding to three standard deviations. Therefore, we can assume that codons for which CEI modulus is greater than 3 have a significant effect on the protein expression level.

To better understand the process of protein synthesis we introduce Codon Productivity (CP) metric. Productivity can be interpreted as the cell’s amino acid production from the specific codon, which is defined by codon and corresponding tRNA frequencies, and can be used to estimate the average expected number of amino acid copies included in the process of protein formation based on a single codon.

The dots in the Figure 5 represent productivity values for codons. The standard deviation calculated from three independent experimental measurements of protein expression is plotted as a bar for each codon. This error is due to the inaccuracy of the experimental measurements. The productivity of most synonymous codons differs by more than the error range, therefore these differences can be considered statistically significant. Codon productivity shows the average number of amino acids produced by a cell per codon.

**Figure 5.**
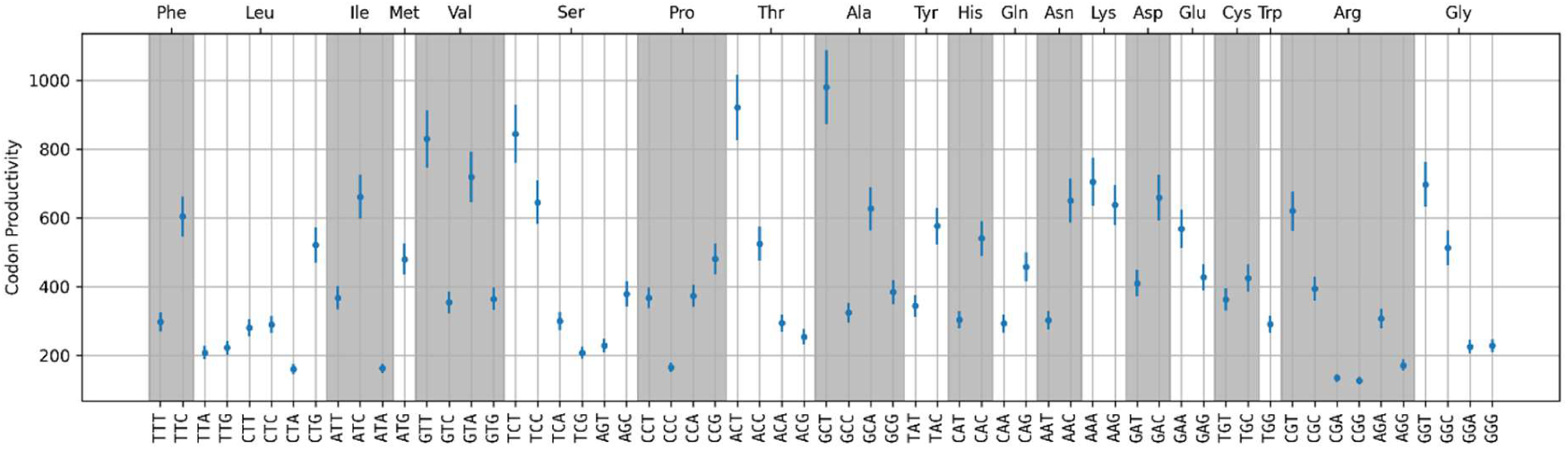
Codon productivity (CP) values show the average amount of amino acids produced by the cell from a single codon. CP values are shown as blue dots. Blue vertical lines are showing the error range corresponding to a single standard deviation. Error estimation is based on the difference between three independent experimental expression measurements.

Codon Productivity (CP) and Codon Expression Index (CEI) values for each codon are listed in the Supplementary Table 1. CP and CEI achieve a very high degree of linear correlation (0.945). That is, both of our proposed metrics can be used interchangeably to analyse the protein expression. The differences between the methods are due to the different nature of the errors: for CEI the main factor is the use of rank correlation, and for codon productivity it is the fact that we calculated it based on a limited set of genes from the organism, those ones for which the expression levels are known.

Interestingly, gene length does not affect its expression level in *E. coli*, unlike higher eukaryotes, for which there is a positive correlation (38). For *E. coli*, the correlation between the length of a gene and the expression level of the corresponding protein is −0.13 based on the used dataset.

Both of the proposed metrics: Codon Expression Index (CEI) and Codon Productivity, were used to predict the expression level of *E. coli* genes based on their nucleotide sequences. Prediction models were trained and tested using 11-fold cross-validation. Both methods achieve r=0.70 linear correlation between the predicted and actual expression values (Figure 6).

**Figure 6.**
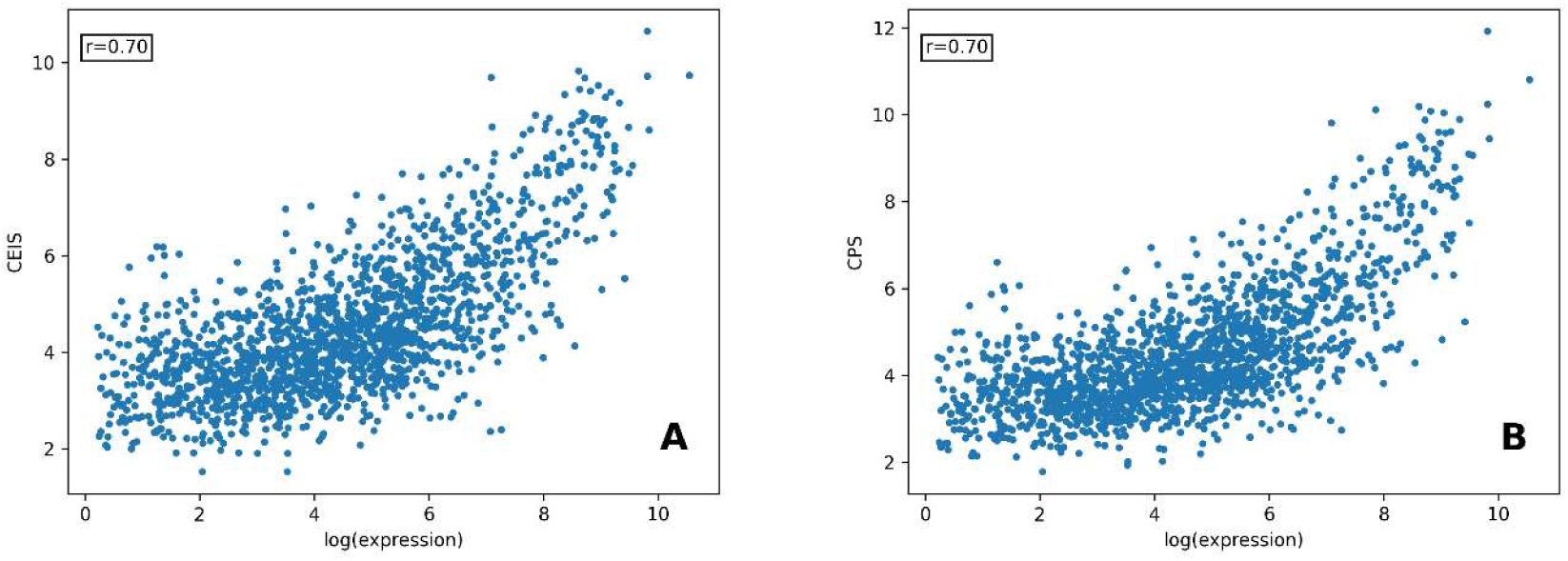
Validation of the expression prediction models on the *E. coli* dataset using 11-fold cross-validation. A. Codon expression index score (CEIS) is based on the CEI values for codon influence on the integral protein expression. B. Codon productivity score (CPS) is based on the CP values for the average number of amino acids produced from a single codon. Both models achieve an r=0.70 linear correlation coefficient between the predicted and actual log expression values.

We compared the prediction accuracy of both models with other existing methods for expression prediction on the same *E. coli* dataset. We tested Codon adaptation index (CAI), tRNA adaptation index (TAI) and Relative codon bias score (RCBS) on disjoint training and test sets. Correlation coefficients between the predicted and experimental log values are r=0.62 for CAI and r=0.54 for TAI (Figure 7). We also tested the RCBS model (10), which has the highest correlation between the predicted and actual expression values, according to the source at r=0.70. When tested on the dataset we used, the efficiency of this method turned out to be significantly lower at r=0.51.

**Figure 7.**
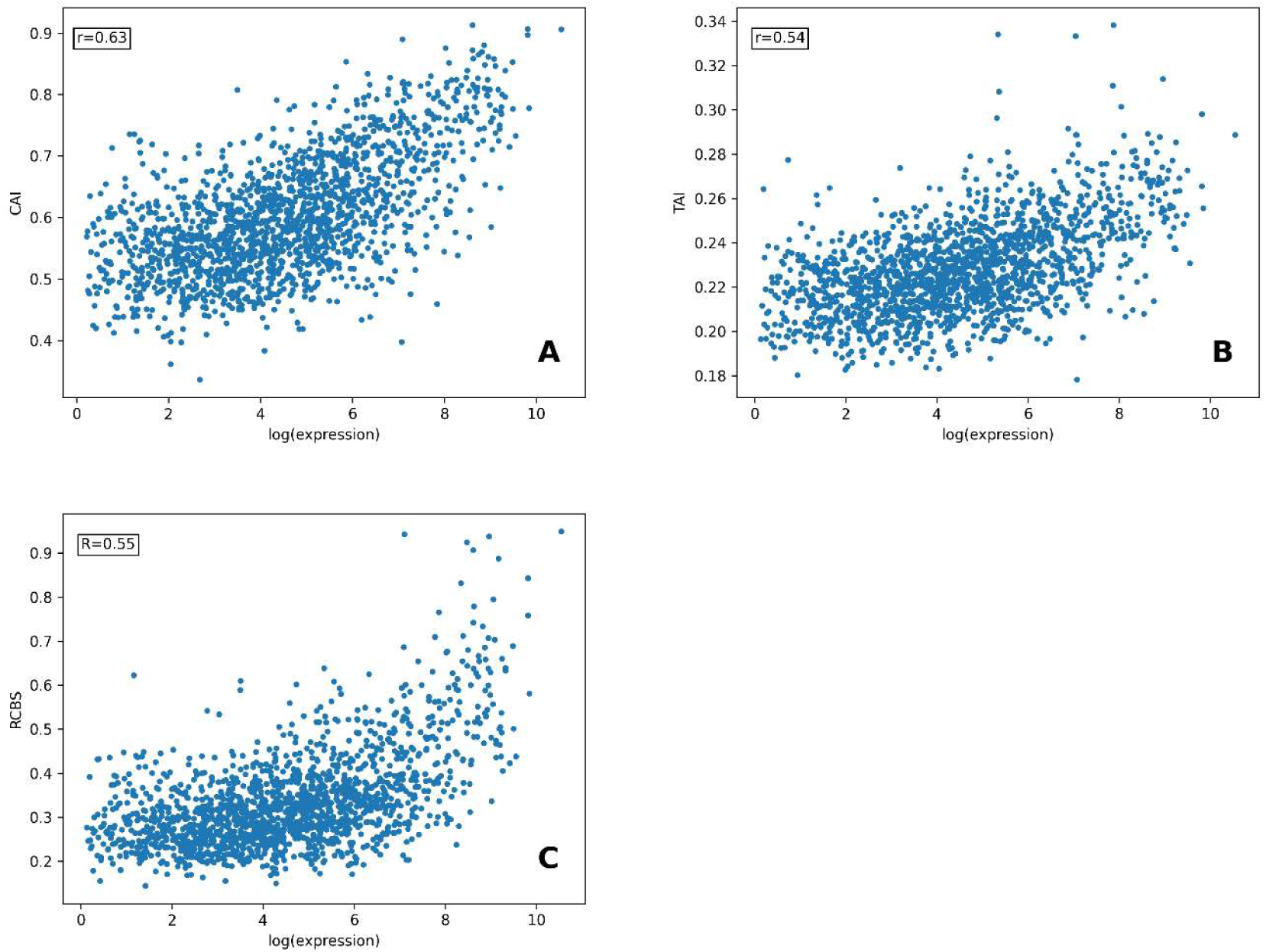
Validation of the models for expression prediction on the *E. coli* dataset. A. Codon Adaptation Index (CAI) achieves an r=0.63 linear correlation coefficient with log expression values. B. tRNA Adaptation Index (TAI) achieves an r=0.54 linear correlation coefficient with log expression values. C. Relative codon bias score (RCBS) achieves an r=0.55 linear correlation coefficient with log expression values.

In order to show the influence of correlations between codons on gene expression, we proposed the Codon Pair Expression Index (similar to the CEI but calculated using codon pair frequencies instead of the individual codon frequencies) and created an expression prediction model for this index. CPEI values are listed in the Supplementary Table 2. Pearson’s linear correlation coefficient between the predicted and actual log expression level is r=0.71 (See Figure 8). The prediction accuracy for CPEIS is moderately higher than for CEIS, while the computational complexity increases significantly, which does not allow us to recommend using this method for expression prediction.

**Figure 8.**
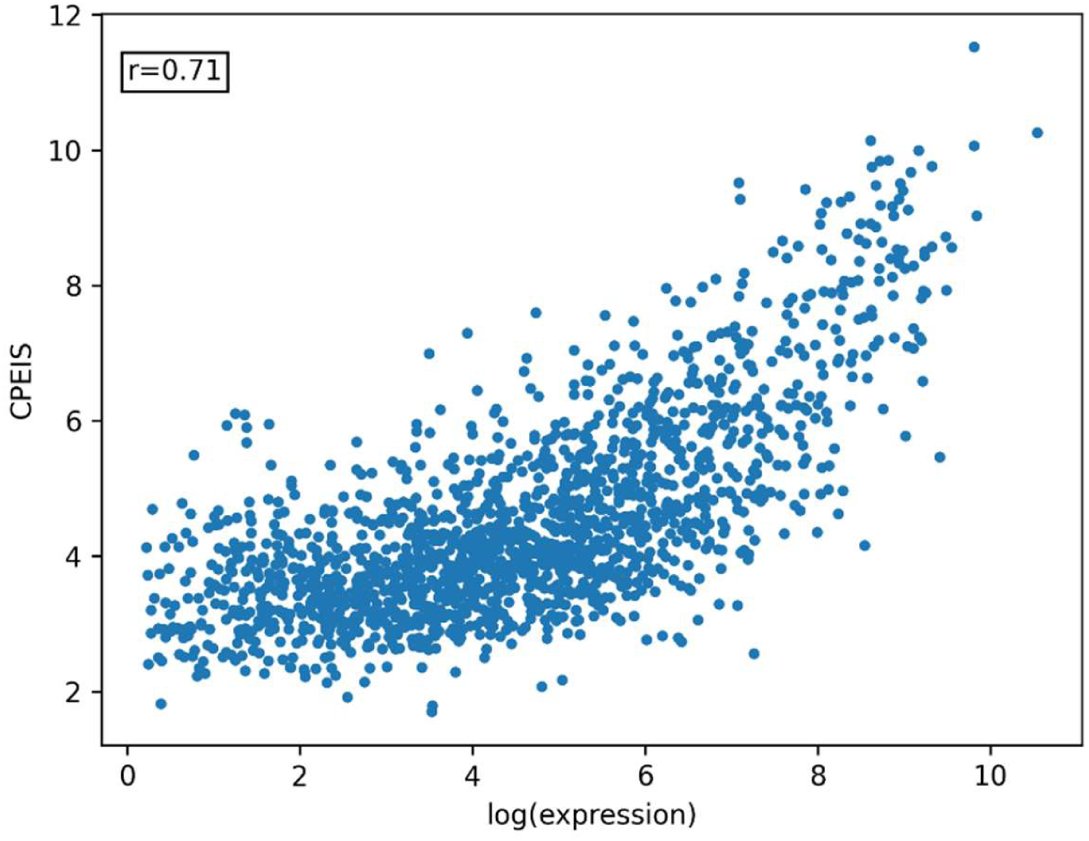
Validation of the expression prediction accuracy for Codon pair expression index score (CPEIS) using 11-fold cross-validation. CPEIS is based on CPEI values for the influence of codon pairs on the integral protein expression. This modal achieves an r=0.71 linear correlation coefficient between the predicted and actual log expression values.

There are 3721 possible codon pairs, that do not contain a stop codon. Genes from the used dataset contain all the pairs except for 14: TCTAGC, CCCCTA, ATTAGG, GTTAGG, TCTAGA, TCTAGG, TCGAGG, CCTAGA, CCTAGG, ACTAGG, GCTAGA, GCTAGG, TATAGG, CGGAGA.

As can be seen from Table 2, for all methods for expression prediction, the peak prediction accuracy is achieved for genes with a log expression level above 2 (more than 6 protein copies per cell). All prediction methods in the table are based on nucleotide sequence analysis; therefore, it can be concluded that the expression regulation for genes with a low expression level is achieved by other means of regulation. At the same time, for genes with higher expression levels, codon frequency is a major regulatory factor. Based on this, and the fact that the codon distributions for four classes of *E. coli* genes with different expression levels are highly similar, we suggest that all genes are optimised for a certain organism regardless of their expression levels.

**Table 2.**
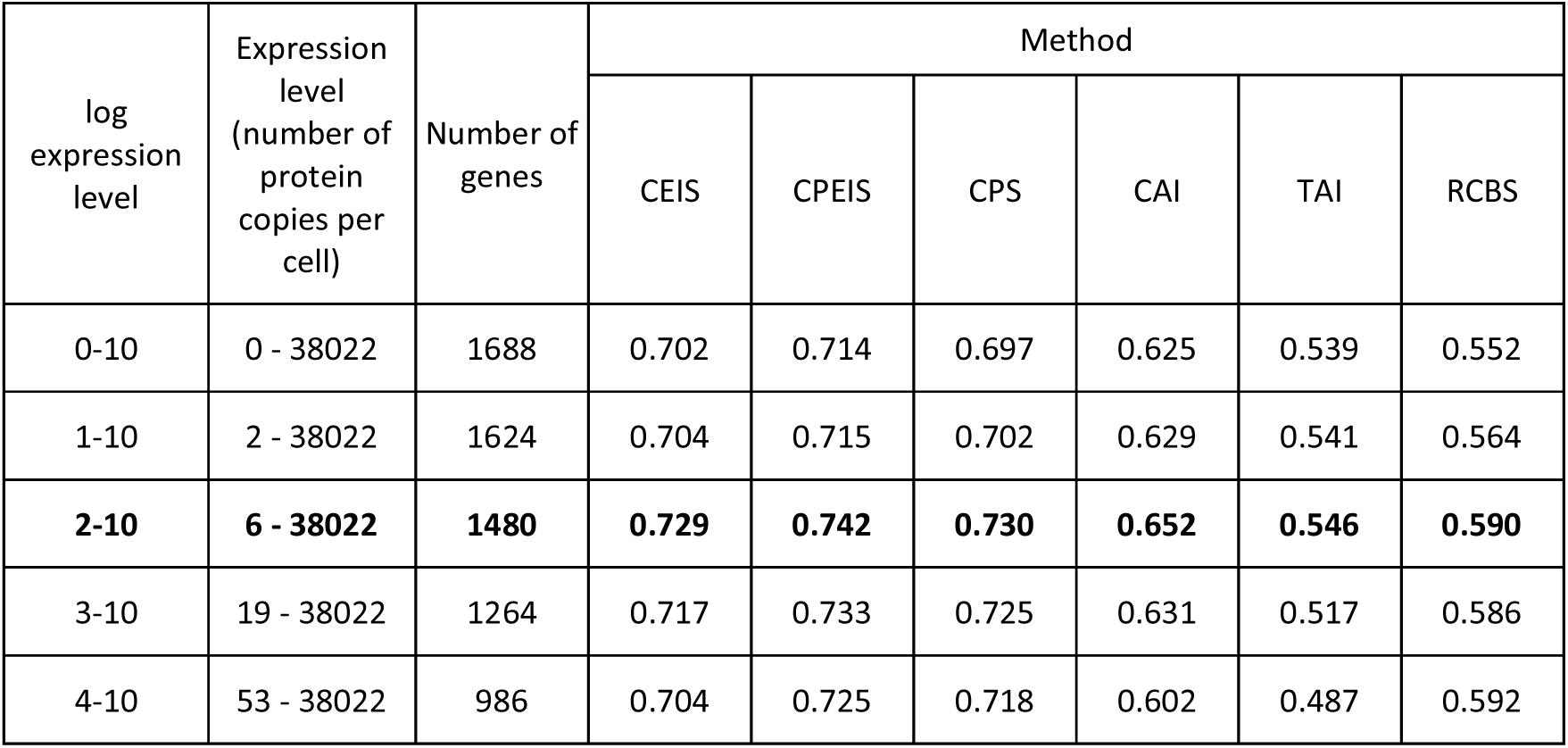
Expression prediction accuracy for *E. coli* genes with different levels of protein expression. Prediction accuracy is measured as a coefficient of linear correlation between the predicted and actual log expression values.

| log expression level | Expression level (number of protein copies per cell) | Number of genes | Method |  |  |  |  |  |
| --- | --- | --- | --- | --- | --- | --- | --- | --- |
|  |  |  | CEIS | CPEIS | CPS | CAI | TAI | RCBS |
| 0-10 | 0 - 38022 | 1688 | 0.702 | 0.714 | 0.697 | 0.625 | 0.539 | 0.552 |
| 1-10 | 2 - 38022 | 1624 | 0.704 | 0.715 | 0.702 | 0.629 | 0.541 | 0.564 |
| **2-10** | **6 - 38022** | **1480** | **0.729** | **0.742** | **0.730** | **0.652** | **0.546** | **0.590** |
| 3-10 | 19 - 38022 | 1264 | 0.717 | 0.733 | 0.725 | 0.631 | 0.517 | 0.586 |
| 4-10 | 53 - 38022 | 986 | 0.704 | 0.725 | 0.718 | 0.602 | 0.487 | 0.592 |

We created a Python module for calculation of the CEI and CP values and for expression prediction based on these indices. Package is available for download from PyPI as “cei”. Source code is available in the Supplementary materials as well as on Github (https://github.com/conzaytsev/CodonExpressionIndex).

## DISCUSSION

Protein expression level is determined by a variety of factors. These include promoter strength, transcription rate, ribosome binding site strength, translation rate, degradation rates of both mRNA and protein, as well as the influence of regulatory elements. The presence of all these factors allows for the same expression level to be achieved through numerous distinct combinations (39). One of the most important parameters affecting protein expression is the frequency of synonymous codon use (1, 25, 40–42). Codon usage primarily impact the speed and efficiency of the translation process (elongation) due to the varying frequencies of the corresponding tRNA molecules. However, several studies have also shown that codon usage can affect the transcription and stability of the mRNA itself (3, 43).

We demonstrated that the frequencies of individual codons correlate with the protein expression level in E. coli. To capture this relationship, we proposed two new metrics. The first is the Codon Expression Index (CEI), which characterises the influence of a specific codon on the overall protein expression level. The second is Codon Productivity (CP), which measures the average number of amino acid copies produced by a cell based on the usage of a particular codon. Interestingly, these two quite different methods - the statistical CEI and the empirical CP - showed a very high mutual correlation, with a correlation coefficient of r=0.945 between their respective values. This suggests the two metrics can be considered interchangeable, though they have distinct interpretations. The CEI indicates the direction and strength of a particular codon’s influence on the integral level of protein expression. In contrast, the codon productivity (CP) value represents the expected number of copies of an amino acid that should be obtained from the use of that specific codon.

We used both of the metrics we developed for protein expression prediction and called the prediction methods: Codon Expression Index Score (CEIS) and Codon Productivity Score (CPS). When we evaluated the performance of these models, we found that the correlation coefficient between the predicted expression levels of E. coli genes and the experimentally measured protein expression levels was 0.7 for both models. This level of correlation represents a significant improvement over all currently widely used predictors. Specifically, our models demonstrated a 13% higher correlation relative to the Codon Adaptation Index (CAI). Other common methods like the tRNA Adaptation Index (TAI) and Ribosome Binding Site Strength (RBCS) were found to be less effective than CAI on the current dataset. The prediction accuracy was nearly identical between our two models, CEIS and CPS. This is not surprising given the high correlation we previously observed between the CEI and CP metrics themselves.

Our protein expression level prediction models offer several advantages over other popular methods, including greater prediction accuracy. Unlike the widely-used Codon Adaptation Index (CAI), our models do not impose artificial limitations, such as restricting each amino acid to the single optimal codon. Instead, CEI calculates the degree of influence each codon has on protein expression, which can be either positive or negative. Additionally, our models can be trained on expression data for any organism under any condition. We discovered that our methods do not always align with CAI regarding preferred codons. For example, according to CAI, the codon ACC is the preferred one for threonine because it is the most common in highly expressed genes (Supplementary Table 1). However, both our CEI and CP methods indicate that the codon ACT is more effective at protein production. Similar discrepancies were found for serine, alanine, and aspartic acid. This difference arises because, unlike CAI, our methods analyse the dynamics of codon frequency changes among genes with varying expression levels, rather than simply identifying the most frequent codon in a set of highly expressed genes.

The primary drawback of our models is that they require experimental data on protein expression for the genes of an organism in question. However, we believe that as proteomic data become more widely available for popular species, these models will be able to provide more accurate practical results.

We also computed a range of CPEI values for codon pairs and developed an expression prediction model based on them. The accuracy of this model showed a slight increase compared to the model based on individual codons. It is worth noting that correlations between neighbouring codons in *E. coli* have been previously documented (44). Experimental evidence has also confirmed that correlations between codons can impact protein expression. For example, using codon pairs has notably enhanced the expression of synthetic sequences (7), while consecutive CGA codons (arginine) were found to disrupt expression in *E. coli* (45). We believe that the marginal improvement in prediction accuracy using codon pairs is due to the primarily negative influence of certain codon combinations. These combinations may already be depleted due to the natural optimisation of genes during the evolution of a specific organism’s genome. However, considering codon pairs when optimising gene sequences for recombinant expression purposes could have a significant impact on the protein expression levels.

It is important to note that for genes with the lowest expression levels (0-6 gene copies per cell), the correlation between predicted and experimental expression values is significantly lower than average. When these genes are excluded from the dataset, the prediction accuracy increases significantly for all considered methods. This leads us to assume that the lowest expression levels are controlled by more subtle mechanisms of expression regulation, rather than by codon frequencies. This might be because a combination of strong transcription with weak translation is not energy efficient (39).

Our results suggest that at least 70% of the protein expression level is determined by codon frequencies, while the remaining regulation, which is not codon related, provides fine-tuning of the expression level.

We have developed our methods based on experimental data on the expression level of *E coli* proteins; therefore, expression prediction using the proposed methods is effective for *E. coli* genes. However, for exogenous sequences (genes from other organisms), the prediction accuracy may become significantly lower. That is because those sequences are not optimised for *E. coli*, and lots of different expression limitations may arise. Despite this potential decrease in prediction accuracy for exogenous sequences, the influence of codon frequencies on protein expression should remain. Therefore, both CEI and CP indices will likely continue to be relevant and useful for assessing protein expression levels.

Despite the differences in the genes associated with adaptation to different expression levels, codon distributions across all classes of *E. coli* genes are much more similar to each other than to the Alien Class consisting of genes originating from other species. This suggests that *E. coli* has a species-specific optimisation of its gene sequences, regardless of their expression level. The goal of this optimisation can be both to achieve the balance between codon usage and tRNA pool ratios, and to avoid the formation of toxic mRNAs (46). The application of our CEI model offers a superior alternative to traditional methods for assessing expression levels within the cellular environment.

## DATA AVAILABILITY

Genomic sequence for *E. coli* strand ATCC 25922 and all the list of genes were derived from a source in the public domain: https://genomes.atcc.org/genomes/ccbc9e61ad334c2c

Expression data for the genes from *E. coli* strand ATCC 25922 was derived from https://doi.org/10.1016/j.dib.2014.08.004

“cei” module for Python is available for download from PyPI or from Github (https://github.com/conzaytsev/CodonExpressionIndex)

Source code for “cei” Python module is available in the Supplementary material.

## SUPPLEMENTARY DATA

**Supplementary Table 1.** This table contains Codon expression index (CEI) values and Codon productivity (CP) values for each of the codons as well as their abundance in the class of *E. coli* genes with the highest protein expression levels.

**Supplementary Table 2.** This table contains Codon pair expression index (CPEI) values for each of the codon pairs.

## ACKNOWLEDGEMENTS

We are grateful to Eugene Korotkov, Maria Yurkova, and Dmitrii Kostenko for their seminal discussion and support.

