## Supplemental Table 1 for "Exploring individual codon influence on protein expression: a predictive approach"

**CEI and CP Summary Table.**

“AA copies per codon” column is the total amount of a certain amino acid produced by the cell from the corresponding codon.
“Codon copies in dataset” column is the total number of a certain codon in the *E. coli* dataset.
“Codon productivity” is calculated as a ratio of “AA copies per codon” to “Codon copies in the dataset”.
“Codon copies in Class 1” is the total number of each of the codons in the class of *E. coli* genes with the highest expression numbers. It is used by the Codon adaptation index to determine the preferred codons.
“Codon expression index” is a measure of codon influence on the integral protein expression level of a gene.

| AA | Codon | AA copies per codon | Codon copies in dataset | Codon copies in Class 1 | Codon expression index | Codon productivity |
|---|---|---|---|---|---|---|
| Phe | TTT | 3392215 | 11371 | 1808 | -15,62 | 298±27 |
| | TTC | 6156987 | 10182 | 2774 | 7,8 | 605±57 |
| Leu | TTA | 1372317 | 6571 | 761 | -17,81 | 209±19 |
| | TTG | 1674723 | 7506 | 1021 | -17,36 | 223±19 |
| | CTT | 1627308 | 5794 | 905 | -12,85 | 281±25 |
| | CTC | 1777978 | 6128 | 1029 | -10,44 | 290±25 |
| | CTA | 299309 | 1856 | 194 | -14,76 | 161±14 |
| | CTG | 17750260 | 34032 | 8005 | 3,86 | 522±51 |
| Ile | ATT | 6405566 | 17394 | 3370 | -7,6 | 368±34 |
| | ATC | 10410008 | 15740 | 4445 | 10,6 | 661±64 |
| | ATA | 215098 | 1318 | 138 | -13,84 | 163±13 |
| Met | ATG | 7773239 | 16178 | 3708 | 0,33 | 480±46 |
| Val | GTT | 9306907 | 11213 | 3522 | 13,08 | 830±84 |
| | GTC | 3121197 | 8786 | 1704 | -7,12 | 355±31 |
| | GTA | 4814578 | 6696 | 1848 | 8,43 | 719±73 |
| | GTG | 5817561 | 15941 | 3188 | -6,42 | 365±33 |
| Ser | TCT | 4262417 | 5046 | 1699 | 11,46 | 845±84 |

| | | | | | | |
|---|---|---|---|---|---|---|
| | TCC | 3429798 | 5314 | 1501 | 6,7 | 645±63 |
| | TCA | 943551 | 3138 | 512 | -9,00 | 301±27 |
| | TCG | 1005401 | 4817 | 671 | -15,01 | 209±18 |
| | AGT | 940612 | 4108 | 539 | -13,85 | 229±20 |
| | AGC | 3519700 | 9275 | 1761 | -4,10 | 379±36 |
| **Pro** | CCT | 1402942 | 3817 | 724 | -4,73 | 368±30 |
| | CCC | 427069 | 2572 | 249 | -17,10 | 166±14 |
| | CCA | 1849491 | 4943 | 1003 | -4,07 | 374±33 |
| | CCG | 7419731 | 15424 | 3588 | 0,78 | 481±44 |
| **Thr** | ACT | 4829225 | 5237 | 1672 | 12,60 | 922±95 |
| | ACC | 7567292 | 14407 | 3532 | 4,62 | 525±50 |
| | ACA | 915339 | 3101 | 492 | -8,51 | 295±25 |
| | ACG | 2010523 | 7877 | 1278 | -12,97 | 255±22 |
| **Ala** | GCT | 9368522 | 9552 | 3050 | 11,44 | 981±108 |
| | GCC | 4806426 | 14765 | 2668 | -10,54 | 326±29 |
| | GCA | 7435500 | 11844 | 3118 | 8,18 | 628±63 |
| | GCG | 8145107 | 21120 | 4449 | -6,14 | 386±35 |
| **Tyr** | TAT | 2926522 | 8484 | 1579 | -9,05 | 345±32 |
| | TAC | 4311932 | 7473 | 1998 | 5,23 | 577±53 |
| **His** | CAT | 2218642 | 7301 | 1175 | -10,82 | 304±25 |
| | CAC | 3420104 | 6321 | 1612 | 3,30 | 541±51 |
| **Gin** | CAA | 2506994 | 8534 | 1337 | -10,86 | 294±27 |
| | CAG | 8285246 | 18063 | 3844 | -3,4 | 459±43 |
| **Asn** | AAT | 2628454 | 8665 | 1468 | -9,25 | 303±27 |
| | AAC | 8912002 | 13684 | 3810 | 12,01 | 651±64 |
| **Lys** | AAA | 14714089 | 20869 | 5970 | 17,04 | 705±71 |
| | AAG | 3823969 | 5993 | 1569 | 6,93 | 638±57 |

| | | | | | | |
|---|---|---|---|---|---|---|
| **Asp** | GAT | 8146115 | 19817 | 4270 | -0,35 | 411±38 |
| | GAC | 8319457 | 12620 | 3570 | 11,99 | 659±66 |
| **Glu** | GAA | 15004733 | 26349 | 6918 | 10,87 | 569±55 |
| | GAG | 4715266 | 11015 | 2355 | -2,26 | 428±39 |
| **Cys** | TGT | 942800 | 2593 | 545 | -6,2 | 364±32 |
| | TGC | 1536165 | 3612 | 763 | -3,82 | 425±40 |
| **Trp** | TGG | 2308168 | 7906 | 1285 | -14,12 | 292±25 |
| **Arg** | CGT | 9257924 | 14914 | 4119 | 11,22 | 621±57 |
| | CGC | 5535695 | 14029 | 2737 | -5,14 | 395±36 |
| | CGA | 211629 | 1564 | 126 | -15,77 | 135±12 |
| | CGG | 311218 | 2438 | 192 | -18,14 | 128±11 |
| | AGA | 164818 | 536 | 68 | -8,7 | 307±29 |
| | AGG | 54611 | 318 | 23 | -10,5 | 172±16 |
| **Gly** | GGT | 11223904 | 16081 | 4703 | 14,96 | 698±65 |
| | GGC | 9459232 | 18401 | 4428 | 4,35 | 514±50 |
| | GGA | 805090 | 3561 | 482 | -13,39 | 226±19 |
| | GGG | 1324437 | 5776 | 833 | -15,15 | 229±19 |
