## Supplemental Table 2 for "Exploring individual codon influence on protein expression: a predictive approach"

**Codon pair expression index values**

Second codon in pair

| First codon in pair | TTT | TTC | TTA | TTG | CTT | CTC | CTA | CTG | ATT | ATC | ATA | ATG | GTT | GTC | GTA | GTG | TCT | TCC | TCA | TCG | AGT | AGC | CCT | CCC | CCA | CCG | ACT | ACC | ACA | ACG | GCT | GCC | GCA | GCG | TAT | TAC | CAT | CAC | CAA | CAG | AAT | AAC | AAA | AAG | GAT | GAC | GAA | GAG | TGT | TGC | TGG | CGT | CGC | CGA | CGG | AGA | AGG | GGT | GGC | GGA | GGG |
|---|---|---|---|---|---|---|---|---|---|---|---|---|---|---|---|---|---|---|---|---|---|---|---|---|---|---|---|---|---|---|---|---|---|---|---|---|---|---|---|---|---|---|---|---|---|---|---|---|---|---|---|---|---|---|---|---|---|---|---|---|---|
| TTT | -6,77 | -7,06 | -8,38 | -9,92 | -6,14 | -6,48 | 0,22 | -6,99 | -6,88 | -3,92 | -4,18 | -3,91 | -1,73 | -7,56 | -0,64 | -3,57 | -3,27 | -4,57 | -5,83 | -6,73 | -3,13 | -4,88 | -3,74 | -4,37 | -2,98 | -7,31 | -1,78 | -6,23 | -2,72 | -5,34 | -3,54 | -6,98 | 1,31 | -4,58 | -5,79 | -4,83 | -8,03 | -4,1 | -3,72 | -6,31 | -5,63 | -3,36 | -0,17 | 0,1 | -6,05 | -0,31 | -2,35 | -2,99 | -3,36 | -4,97 | -3,36 | -6,31 | -7,47 | -2,85 | -2,14 | 0,27 | 0,04 | -2,91 | -7,22 | -1,35 | -2,89 |
| TTC | -3,43 | 1,96 | -1,76 | -2,28 | -2,09 | 1,08 | 1,25 | 2,31 | 0,01 | 3,26 | -1,37 | 4,29 | 5,6 | -0,06 | 3,83 | 1 | 2,82 | 0,91 | -1,85 | -4,2 | -4,12 | -0,64 | -0,33 | -4,59 | -0,87 | 2,23 | 1,73 | 0,98 | -1,08 | -2,17 | 0,87 | -4,27 | 0,47 | -0,14 | -1,07 | 1,87 | -1,22 | 1,16 | 0,38 | 2,14 | -2,18 | 4,62 | 6,66 | 1,37 | -0,69 | 2,93 | 4,37 | 0,2 | -1,53 | 0,53 | -6,63 | 4,49 | 0,64 | -1,98 | -1,91 | 0,87 | -3,09 | 3,84 | 0,64 | 0,44 | -0,6 |
| TTA | -6,64 | -4,18 | -6,67 | -9,5 | -4,66 | -5,16 | -3,91 | -8,21 | -5,66 | -7,25 | -4,47 | -7,31 | -2,42 | -1,95 | 1,3 | -6,08 | -4,2 | -1,78 | -3,47 | -6,06 | -7 | -6,56 | -4,92 | -7,44 | -2,43 | -7,03 | -0,56 | -4,93 | -3,76 | -6,52 | -0,66 | -3,82 | -2,56 | -5,91 | -4,74 | -2,56 | -4,18 | -4,07 | -5,62 | -6,52 | -6,89 | -4,11 | -4,19 | -2,83 | -0,97 | 0 | -2,09 | -0,92 | -5,47 | -3,4 | -7,68 | -6,06 | -7,02 | -2,18 | -3,24 | -2,38 | -1,45 | -4,98 | -3,36 | -2,85 | -4,78 |
| TTG | -7,11 | -4,21 | -7,01 | -6,87 | -6,49 | -7,27 | -6,54 | -10,64 | -8,37 | -3,86 | -4,31 | -5,8 | -1,24 | -4,34 | -1,32 | -7,28 | -0,76 | -2,38 | -3,26 | -5,82 | -5,11 | -3,93 | -3,48 | -7,52 | -4,72 | -8,46 | -0,13 | 1,47 | -2,32 | -4,5 | -1,28 | -3,26 | -3,95 | -4,54 | -2,6 | -3,21 | -5,83 | -2,41 | -7,34 | -5,2 | -4,16 | -0,7 | -1,89 | -2,07 | -2,24 | -1,95 | -1,04 | -2,58 | -2,08 | -3,56 | -5,86 | -4,24 | -5,94 | -2,76 | -5,21 | -0,29 | 0,15 | -0,22 | -0,41 | -2,77 | -6,72 |
| CTT | -4,66 | -4,96 | -4,84 | -5,38 | -1,27 | -0,4 | -1,91 | -1,2 | -6 | -4,05 | -2,25 | -1,99 | -0,68 | -3,36 | -0,01 | -3,17 | -3,56 | -3,95 | -2,92 | -5,79 | -4,09 | -4,79 | -3,98 | -4,75 | -3,05 | -2,1 | -3,41 | -3,71 | -0,16 | -3,15 | -2,09 | -5,57 | -1,7 | -3,88 | -6,76 | -4,13 | -4,18 | -2,34 | -0,05 | -5,5 | -5,51 | -3,86 | -3,34 | -3,21 | -5,62 | 1,31 | -4,84 | -2,09 | -5,11 | -2,21 | -3,6 | -0,35 | -2,71 | -3,35 | -1,96 | -1,6 | 0 | -2,05 | -2,58 | -2,25 | -3,17 |
| CTC | -5,94 | -5,13 | -5,22 | -1,68 | -2,39 | -1,46 | -2,13 | -3,43 | -5,15 | -1,22 | -0,45 | -1,15 | -1,2 | -2,18 | 0,5 | -0,39 | 0,85 | -1,68 | -4,22 | -4,56 | -3,31 | -4,2 | -1,6 | -3,95 | -1,38 | -1,61 | -2,53 | -4,2 | -0,6 | -5,05 | -2,36 | -8,29 | -0,35 | -5,3 | -4,67 | -2,83 | -3,84 | -3,35 | -1,44 | -1,78 | -6,38 | -3,77 | -3 | -0,94 | -4,13 | 0 | -0,07 | -0,28 | -0,67 | -4,66 | -7,98 | 0,06 | -1,63 | -2,03 | -2,4 | -0,96 | 0,4 | -1,88 | -2,49 | -2,56 | -4,03 |
| CTA | -5,88 | -3,84 | -5,87 | -5,87 | -2,78 | -1,31 | -3,07 | -5,93 | -4,7 | -0,88 | -1,56 | -3,08 | 2,26 | -0,09 | 0,68 | -0,14 | -3,53 | -2,04 | -2,67 | -4,34 | -4,68 | -2,43 | -1,71 | -1,66 | -3,51 | -3,87 | 0,2 | -3,65 | -2,05 | -4,29 | -0,85 | -2,44 | 0,32 | -2,93 | -4,01 | -0,75 | -3,89 | -2,94 | -5,31 | -4,24 | -2,62 | -2,86 | -4,87 | -3,64 | -4,38 | 0,45 | -0,87 | -2,34 | -1,99 | -0,16 | -3,54 | -2,96 | -4,94 | -2,41 | -3,41 | 0,12 | 0,51 | -0,97 | -1,3 | -1,73 | -2,37 |
| CTG | -8,75 | -0,44 | -6,72 | -2,95 | -7,16 | -4,15 | -4,48 | 1,98 | -2,8 | 1,46 | -6,86 | -1,37 | 3,59 | -4,64 | 0,79 | -8,01 | 8,73 | 5,63 | -3,53 | -4,45 | -2,83 | 0,92 | -1,93 | -6,27 | -0,15 | 2,35 | 4,04 | 2,14 | -5,52 | -4,47 | 4,14 | -4,28 | -1,27 | -7,74 | -0,78 | 4,59 | -3,33 | 2,1 | -5,76 | 3,86 | -3,25 | 4,52 | 6,63 | -0,29 | -0,07 | 3,4 | 2,52 | -3,41 | -2,56 | -0,68 | -3,47 | 4,52 | -1,58 | -5,55 | -6,68 | 1,38 | -1,97 | 5,06 | 1,91 | -5,07 | -6,07 |
| ATT | -5,58 | -3,87 | -7,6 | -9,59 | -3,05 | -4,72 | -2,66 | -1,91 | -7,51 | -2,9 | -2,3 | -3,85 | -3,65 | -6,11 | -3,22 | -6,23 | -0,37 | -1,63 | -3,74 | -4,43 | -3,69 | -4,38 | -3,34 | -5,66 | -2,18 | -2,45 | -0,04 | -4,65 | -0,5 | -2,33 | -0,01 | -7,43 | -0,46 | -4,41 | -3,15 | -3,13 | -3,56 | 1,61 | -2,15 | -2,1 | -4,77 | -1,32 | -0,57 | 2,27 | -1,2 | 0,04 | 0,7 | -3,64 | -0,45 | -0,59 | -1,23 | 0,97 | -1,76 | -2,45 | -2,77 | 1,58 | 0,33 | -2,51 | -5,88 | -1,22 | -2,29 |
| ATC | -1,88 | 1,51 | -3,93 | -4,43 | -2,73 | -0,42 | -1,83 | 1,33 | 0 | 1,36 | -1,01 | 0,81 | 8,05 | 0,41 | 0,71 | 0,95 | 3,4 | 2,73 | -1,77 | -1,55 | -4,67 | -3,98 | 1,01 | -3,91 | -0,74 | 2,2 | 3,4 | 0,59 | -2,02 | -3,76 | 6,15 | -4,34 | 2,16 | 2,02 | 0,79 | 0,45 | -1,27 | -0,72 | -0,72 | -0,51 | -3,36 | 2,22 | 5,08 | 2,88 | 1,85 | 3,55 | 7,17 | 3,19 | -0,93 | -0,67 | -4,94 | 6,52 | -1,78 | -0,57 | -3,61 | 0,67 | 0,19 | 5,29 | 0,66 | -0,06 | -3,51 |
| ATA | -5,06 | -3,24 | -4,5 | -5,06 | -3,22 | -1,65 | 0,2 | -3,14 | -2,86 | -1,58 | -1,25 | -2,8 | -1,03 | -2,64 | 0,45 | -2,28 | -1,8 | -0,69 | -2,22 | -2,9 | -2,73 | -3,64 | -2,77 | -2,02 | -1,47 | -3,68 | -2,18 | -5,26 | -1,91 | -3,24 | -1,21 | -3,62 | -3,39 | -3,59 | -3,21 | 0,33 | -2,14 | -1,19 | -2,61 | -1,62 | -3,45 | -2,78 | -4,04 | -2,45 | -3,72 | -2,17 | -4,01 | -4,43 | -1,54 | -0,24 | -4,13 | 0,76 | -2,96 | -0,98 | -0,51 | -2,41 | -1,16 | -1,84 | -3,62 | -3,61 | -3,99 |
| ATG | -4,04 | -1,09 | -5,58 | -7,61 | -6,46 | -4,98 | -2,26 | -2,15 | -3,3 | -1,34 | -4,75 | -2,5 | 0,94 | -2,87 | 1,21 | -4,95 | 0,26 | 1,22 | 1,75 | -4,48 | -4,67 | -0,55 | -2,35 | -4,33 | -1,24 | -3 | 1,24 | -1,38 | -1,7 | -4,12 | 2,96 | -1,57 | 2,17 | -3,55 | -2,38 | 0,64 | -3,48 | 1,19 | -1,48 | -2,19 | -3,37 | 2,81 | 3,9 | 0,92 | 0,19 | 3,01 | 2,72 | -1,47 | 1,82 | -4,21 | -2,39 | 1,97 | -2,99 | -0,25 | -3,76 | -1,74 | -2,19 | 1,7 | 0,01 | -3,39 | -4,95 |
| GTT | -5,52 | -0,42 | -2,32 | -3,7 | 1,2 | 0,99 | -1,53 | 5,94 | -0,93 | 5,2 | -1,59 | 3,45 | 6,74 | 0,95 | 2 | 1,88 | 4,6 | 0,75 | -1,01 | -2,76 | -0,77 | 1,01 | 0,14 | -3,92 | 1,91 | 3,53 | 2,44 | 0,82 | -1,31 | -1,01 | 6,86 | -1,38 | 4,57 | -0,24 | -3,62 | 1,76 | -1,14 | 3,71 | -0,49 | 1,15 | -2,13 | 4,21 | 5,69 | 3,52 | 3,61 | 6,86 | 5,47 | 1,47 | -0,83 | 0,19 | -0,41 | 8,02 | 1,29 | -1,13 | -2,27 | -0,85 | 0 | 2,59 | 1,33 | -1,74 | -2,57 |
| GTC | -4,18 | 1,13 | -5,78 | -2,96 | -2,35 | -1,81 | 0 | -0,77 | -4,16 | -1,24 | -1,61 | -2,28 | 0,46 | -2,07 | 0,89 | -1,05 | 0,1 | -1,37 | -1,77 | -3,16 | -6,92 | -6,06 | 1,09 | -2,35 | -0,7 | -0,27 | -1,13 | -2,19 | -2,59 | -5,75 | 0,73 | -5,9 | -3,19 | -3 | -3,39 | -0,84 | -2,91 | -1,87 | -0,26 | -0,7 | -5,31 | -0,34 | -1,29 | 1,13 | -3,71 | 0,08 | -0,49 | -4,03 | -3,35 | -2,3 | -7,83 | 1,29 | -1,34 | -1,06 | -3,59 | -3,15 | -3,34 | 0,3 | -2,07 | -3,5 | -5,66 |
| GTA | -3,81 | 0,23 | -3,64 | -1,7 | -3,57 | -3,05 | -2,17 | -0,3 | 2,02 | 4,07 | -0,07 | 0 | 5,73 | -1,59 | 1,36 | 1,07 | 3,75 | 2,56 | -1,6 | -2,44 | -2,35 | -0,34 | 1,32 | -2,45 | 0,71 | 0,84 | 5,88 | 1,29 | -0,19 | 0,85 | 4,95 | 0,13 | -0,15 | 1,16 | -0,12 | 2,76 | -1,21 | 1,97 | -0,43 | 1,24 | -0,73 | 5,4 | 0,87 | -3,67 | 1,48 | 4,44 | 4,14 | -1,02 | -0,06 | 1,06 | -0,53 | 3,06 | -1,56 | -3,11 | -3,11 | -0,72 | -1,38 | 6,02 | -1,12 | -1,8 | -3,21 |
| GTG | -4,58 | 0,69 | -4,36 | -6,56 | -2,53 | -6,45 | -3,71 | -4,94 | -2,51 | -0,32 | -3,45 | -5,04 | 3,56 | -2,87 | -3,08 | -7,58 | 1,25 | -0,09 | -2,46 | -4,52 | -3,72 | -1,07 | -1,13 | -4,76 | -0,39 | -4,68 | 4,23 | 1,6 | -3,83 | -5,45 | -0,07 | -2,53 | -2,73 | -7,44 | -1,78 | 0,54 | -3,79 | -1,17 | -4,46 | -5,56 | -2,24 | 0,8 | 4,88 | 0,36 | -1,5 | 0,04 | -3,18 | -4,03 | -3,62 | -1,84 | -3,88 | 0,68 | -4,5 | -4,19 | -5,33 | -4,19 | -2,6 | 1,26 | -0,44 | -5,58 | -5,83 |
| TCT | -0,71 | 3,4 | -5,28 | -1,83 | -0,56 | -0,83 | -2,14 | 3,5 | -2,09 | 2,65 | -0,73 | 3,59 | 2,7 | 0,43 | 3,41 | 1,53 | 4,36 | 1,07 | -1,62 | -2,16 | 0 | -0,13 | -1,29 | -2,87 | 0,32 | 3,96 | 5,44 | 2,94 | 0,03 | -0,29 | 5,6 | -0,93 | 4,94 | 0,64 | -0,01 | 1,35 | -0,85 | 3,09 | 0,42 | 4,61 | -0,91 | 4,37 | 5,85 | 3,17 | 1,53 | 6,37 | 6,73 | 2,5 | 2,35 | -1,07 | -0,91 | 3,04 | 1,92 | -0,99 | -2,39 | 0 | 0 | 4,95 | 1,49 | -1,89 | -1,42 |
| TCC | -3,84 | 1,79 | -3,36 | -0,52 | -0,4 | -1,45 | -0,52 | 2,17 | 0,61 | 4,27 | -2,3 | 2,4 | 2,73 | -1,1 | 2,74 | -0,18 | 1,33 | 1,65 | -2,66 | -2,3 | -2,24 | -1,71 | -0,92 | -4,66 | -0,72 | 1,44 | 0,47 | -0,46 | -1,51 | -0,86 | 4,36 | -0,57 | 2,78 | -0,22 | 0,56 | 3 | 0,14 | 1,1 | -3,26 | 0,27 | -0,2 | 4,35 | 3,77 | 6,15 | 1,32 | 1,52 | 2,17 | 2,41 | -1,61 | 2,48 | -0,04 | 2,2 | -1,33 | -1,66 | -3,99 | -0,95 | -1,19 | 1,86 | -2,13 | -2,02 | -2,18 |
| TCA | -3,89 | -2,35 | -2,77 | -5,15 | -3,68 | -2,96 | -2,63 | -1,69 | -3,71 | -1,82 | -3,16 | -2,72 | -0,79 | -2,63 | -1,55 | -1,21 | 1 | -1,77 | -3,86 | -1,28 | -0,19 | -2 | -2,51 | -3,66 | -3,87 | -2,9 | -0,24 | -0,15 | -3,15 | -3,77 | 1,27 | -4,8 | -2,27 | -4,21 | -2,03 | 0,58 | 0,03 | -1,68 | -3,19 | -0,74 | -3,02 | -0,24 | -0,5 | -1,76 | -0,44 | -1,34 | -0,45 | 0,29 | -1,62 | -3,75 | -4,63 | -2,68 | -3,47 | -3,22 | -3,59 | -1,14 | -1,61 | -0,92 | -0,35 | -2,4 | -2,06 |
| TCG | -5,43 | -1,44 | -4,82 | -5,76 | -4,99 | -3,83 | -4,91 | -7,78 | -5,85 | -2,22 | -3,31 | -5,6 | -3,2 | -2,87 | -2,83 | -5,54 | -1,25 | 0,13 | 0,19 | -3,48 | -0,6 | -1,91 | -2,24 | -4,79 | -3,23 | -4,82 | 0,57 | -1,9 | -0,52 | -4,81 | -1,02 | -3,99 | -2,99 | -7,24 | -2,35 | -0,01 | -4,58 | 0,21 | -5,35 | -1,3 | -4,44 | -2,26 | -0,96 | -0,67 | -3,22 | -1,1 | -3,34 | -3,03 | -2,92 | -0,94 | -4,92 | -4,14 | -5,56 | -3,06 | -4,65 | -0,76 | 0 | -0,23 | -2,78 | -2,82 | -1,97 |
| AGT | -7,83 | -3,78 | -4,45 | -6,12 | -1,89 | -1,21 | -1,91 | -6,54 | -2,57 | -4,8 | -3,07 | -5,91 | -4,4 | -3,05 | -1,5 | -2,95 | -1,08 | -1,3 | -2,77 | -3,87 | -2,61 | -4,57 | -2,34 | -1,8 | -0,56 | -1,45 | -0,7 | -4,88 | -2,59 | -4,35 | -2,01 | -2,8 | 0,18 | -4,85 | -4,25 | -1,72 | -3,67 | -4,41 | -3,85 | -6,11 | -4,47 | -1,13 | -2,86 | -1,29 | -4,49 | -4,1 | -5,17 | -3,7 | -2,66 | -3,53 | -5,29 | -3,23 | -5,07 | -2,48 | -2,59 | -0,26 | 0,51 | -1,52 | -2,6 | -3,61 | -2,92 |
| AGC | -3,52 | 4,1 | -1,19 | -1,36 | -2,86 | -3,2 | 0,13 | 0,27 | -1,73 | 0,82 | -0,4 | -1,14 | -0,49 | -3,31 | 0,34 | -5,13 | 0,57 | 2,08 | 0,42 | -3,33 | -4,87 | -0,93 | -1,9 | -1,98 | -0,22 | -0,99 | 0,35 | -1,92 | -1,43 | -3,62 | -1,54 | -7,27 | 0,31 | -1,88 | -4,36 | -0,32 | -4,25 | -1,54 | -3,51 | -4,51 | -1,33 | -0,75 | -0,53 | 0,4 | -3,31 | -0,61 | -1,89 | -2,75 | -0,54 | -2,07 | -6,71 | 0,65 | -3,88 | -1,55 | -2,39 | 0,95 | -1,23 | -2,25 | -4,45 | -4,01 | -5,04 |
| CCT | -3,17 | -0,24 | -4,69 | -3,72 | -3,37 | -3,58 | -2,41 | -1,96 | -2,74 | -0,6 | -1,24 | -0,88 | -2,11 | -1,84 | -3,52 | -2,76 | 0,64 | -1,99 | -1,85 | -3,17 | -1,41 | 1,15 | -1,41 | -2,31 | -2,39 | 1,76 | 0,17 | -0,97 | -1,41 | -4 | 0 | -4,26 | 1,73 | -2,83 | -1,58 | 0,12 | -3,59 | 0,02 | -1,65 | -1,33 | -2,35 | 0,72 | -1,03 | -0,13 | -2,59 | -1,95 | -2,06 | 0,48 | -2,43 | -1,07 | -0,39 | -1,5 | -1,02 | -1,87 | -0,8 | 0 | 0 | 0,42 | -1,03 | -3,3 | -1,45 |
| CCC | -5,13 | -2,64 | -5,74 | -3,75 | -3,52 | -1,82 | 0 | -1,06 | -5,94 | -1,85 | -0,89 | -2,93 | -3,83 | -3,56 | -3,41 | -6,91 | -3,66 | -2,85 | -1,05 | -3,57 | -1,99 | -3,63 | -1,7 | -0,28 | -2,05 | 0,05 | -1,79 | -4,19 | -4,27 | -5,24 | -2,75 | -6,81 | -4 | -6,03 | -6,32 | -2,43 | -2,22 | -1,72 | -2,54 | -2,73 | -2,89 | -3,58 | -2,88 | -0,26 | -5,46 | -5,6 | -3,8 | -3,35 | -2,65 | -3,38 | -7,11 | -3,38 | -3,86 | -2,22 | -0,8 | -1,83 | -2,17 | -4,87 | -5,4 | -5,08 | -1,47 |
| CCA | -2,21 | -0,99 | -5,79 | -3,87 | -3,54 | -2,08 | -3,25 | -5,62 | -0,79 | 2,89 | -2,06 | -0,22 | 1,16 | -1,23 | 0,24 | -2,98 | -0,78 | 0,33 | -1,67 | -2,39 | -0,41 | 3,18 | -2,14 | -2,62 | -2,14 | -3,23 | 1,31 | 1,62 | -1,89 | -2,75 | 3,35 | -0,79 | -0,13 | -0,95 | -0,83 | 0,75 | -2,21 | -3,4 | -6,68 | -3,36 | -0,55 | 0,55 | -1,49 | -3,56 | -2,91 | -2,53 | -0,5 | -3,14 | 0,91 | 1,89 | -2,17 | 1,04 | -3,82 | -4,8 | -3,26 | 0,22 | -2,64 | -0,06 | -0,32 | -1,17 | -2,23 |
| CCG | -5,92 | 3,7 | -5,62 | -6,26 | -4,67 | -5,12 | -6,67 | -3,34 | -3 | 3,46 | -3,36 | -0,28 | 2,96 | -3,33 | 0,51 | -4,33 | 3,63 | 2,66 | -2,56 | -4,03 | -0,41 | 0,93 | -3,64 | -3,47 | -3,3 | -2,9 | 3,7 | 4,06 | 0,63 | -4,9 | 5,31 | -0,45 | 2,85 | -4,26 | -1,08 | 4,13 | -3,27 | 0,38 | -5,38 | -3,36 | -0,79 | 4,19 | 3,24 | 1,09 | -0,21 | 2,11 | 0,37 | -5,33 | -0,47 | -0,89 | -4,94 | 1,62 | -3,01 | -5,11 | -3,24 | -2,45 | -2,07 | 0,99 | -0,24 | -1,29 | -2,97 |
| ACT | -1,8 | 2,86 | -2,73 | -1,53 | 0,19 | 1,79 | -1,47 | 6,49 | 0,62 | 2,73 | 0,89 | 0,43 | 4,17 | -2,17 | 1,18 | 0,76 | 3,3 | 4,02 | -0,67 | 0,45 | 1,17 | -0,09 | 0,57 | -1,15 | 1,4 | 6,78 | 0,69 | -1,34 | -1,34 | -0,77 | 4,62 | -0,33 | 3,19 | 0,55 | 0,92 | 3,82 | -4,03 | 3,21 | 0,57 | 0,46 | -1,1 | 2,47 | 2,18 | 5,4 | -0,03 | 3,07 | 3,31 | 2,42 | -1,36 | -0,08 | -0,03 | 5,8 | -0,68 | -2,13 | -1,68 | 0,03 | 0 | 3,66 | 4,91 | -2,05 | -0,97 |
| ACC | -3,63 | 1,11 | -4,17 | -4,02 | -2,94 | -2,33 | -1,62 | 3,75 | -2,24 | 2,45 | -2,81 | 1,67 | 4,68 | -4,78 | 5,54 | -0,07 | 3,45 | -0,71 | -2,6 | -4,86 | -3,1 | -2,68 | -3,17 | -1,69 | 0,01 | 1,51 | 1,1 | -0,95 | -2,08 | -4,19 | 0,59 | -5,7 | 0,18 | -2,78 | -1,64 | 0,95 | -2,58 | -1,94 | -1,63 | -0,82 | -1,52 | 4,04 | 5,37 | 4,93 | -2,14 | 2,73 | 1,34 | -2,07 | -0,17 | -2,34 | -3,41 | 2,21 | -1,72 | -5,04 | -2,48 | -1,01 | 1,5 | 7,05 | -2,56 | -3,41 | -4,13 |
| ACA | -3,13 | -1,62 | -7,23 | -6,94 | -2,58 | -3,99 | -3,24 | -4,19 | -2,83 | -0,17 | -2,85 | -1,13 | 0,75 | -0,76 | -0,49 | -2,28 | -0,27 | -0,67 | -2,47 | -3,45 | 0,2 | -0,47 | -0,13 | -1,73 | -2,31 | -3,41 | -2,15 | -0,57 | -1,64 | -3,9 | 0,91 | -0,99 | -1,44 | -1,36 | 0,14 | -1,1 | -3,43 | -3,42 | -4,2 | -2,64 | -2,28 | -1,24 | -1,27 | -2,97 | -4,12 | -1,09 | -1,71 | -2,43 | -0,16 | -3,65 | -2,75 | -0,63 | -2,23 | -2,42 | -0,92 | -1,57 | -1,73 | 0,38 | -1,42 | -2,76 | -2,88 |
| ACG | -4,6 | -2,07 | -7,34 | -4,53 | -6,21 | -5,59 | -6,04 | -4,62 | -2,72 | -2,73 | -3,45 | -4,47 | -2,35 | -2,48 | -2,74 | -4,51 | 0,83 | -0,38 | -2,4 | -2 | -1,67 | -1,39 | -2,33 | -5,61 | -5,23 | -5,59 | -1,69 | -3,11 | -2,7 | -3,37 | -2,52 | -2,19 | -3,85 | -4,97 | -1,17 | -0,6 | -4,24 | -1,21 | -6,06 | -4,88 | -2,56 | 3 | -1,63 | 1,14 | -2,54 | -1,53 | -1,18 | -4,2 | -0,67 | 0,28 | -4,13 | -3,69 | -3,84 | -6,34 | -6,06 | -1,46 | -3,2 | 0,97 | -2,51 | -3,47 | -4,43 |
| GCT | -6,97 | -1,06 | -3,18 | -3,75 | -0,06 | -1,31 | -2,54 | 3,66 | -2,62 | 3,85 | -3,36 | 1,35 | 5,5 | 0,86 | 3,27 | -0,4 | 5,45 | 2,32 | -2,56 | -2,27 | -1,5 | -0,14 | 2,02 | -1,59 | -0,95 | 1,2 | 2,31 | 0,88 | -1,74 | -0,74 | 7,44 | -0,61 | 4,41 | 0,12 | -0,91 | 4,31 | -3,26 | 1,39 | -0,28 | 1,9 | -1,05 | 5,43 | 8,44 | 5,55 | 2,96 | 5,04 | 7,2 | 1,82 | 1,75 | 2,31 | -1,06 | 3,94 | 1,93 | -0,66 | -2,56 | -1,57 | -1,23 | 5,57 | 5,37 | -2,77 | -1,05 |
| GCC | -7,26 | -1,56 | -6,13 | -4,45 | -2,93 | -4,81 | -0,2 | -2,31 | -5,49 | -1,6 | -4,7 | -4,06 | -0,9 | -6,76 | -0,13 | -4,86 | 1,87 | 1,13 | -4,89 | -5,98 | -7,08 | -4,74 | -2,6 | -4,03 | -4,4 | -2,94 | -3,53 | -3,27 | -1,98 | -7,24 | 1,15 | -8,9 | -1,53 | -4,55 | -1,26 | -0,56 | -3,01 | -1,85 | -2,74 | -1,43 | -3,32 | -2,2 | 0,08 | 1,58 | -4,89 | -1,28 | -2,96 | -2,05 | -2,85 | -2,1 | -8,19 | -2,15 | -5,13 | -3,87 | -4,09 | -3,31 | -4,41 | 2,71 | 1,16 | -7,62 | -8,74 |
| GCA | -4,59 | 4 | -4,61 | -3,82 | -3,63 | -2,16 | -2,56 | 1,73 | -0,9 | 1,76 | -2,5 | 1,09 | 1,92 | -3,35 | 1,08 | 0,13 | 2,4 | 2,82 | 0,41 | -3,48 | -2,33 | 0,65 | 2,63 | -1,58 | -0,75 | 2,01 | 4,19 | 1 | 0,1 | -2,63 | 6,52 | -0,44 | 2,05 | 1,89 | 0,96 | 2,77 | -2,07 | 1,02 | -4,26 | -0,97 | -3,78 | 3,32 | 3,39 | -3,34 | -1,34 | 3,84 | 3,25 | -1,5 | 1,18 | -1,3 | -0,72 | 6,26 | -1,65 | -3,56 | -3,29 | -1,08 | -3,55 | 4,13 | 1,11 | -1,46 | -2,22 |
| GCG | -6,8 | 2,15 | -6,53 | -6,48 | -4,93 | -6,54 | -6,53 | -3,14 | -4,25 | 0,52 | -6,86 | -4,82 | 2,07 | -4,2 | -1,97 | -7,47 | 2,71 | 2,14 | -2,79 | -4,08 | -2,41 | -0,24 | -2,6 | -4,3 | -4,31 | -3 | 3,23 | 1,47 | -4,34 | -6,05 | -0,88 | -1,73 | -1,19 | -5,48 | -2,56 | 1,06 | -2,86 | 0,12 | -6,44 | -3,22 | -3 | 2,8 | 4,42 | 0,87 | -2,6 | -0,86 | 0,67 | -4,67 | 0,83 | 1,35 | -5,91 | 0,05 | -1,62 | -6,87 | -6,94 | -3,65 | -2,74 | 3,35 | -1,55 | -4,81 | -6,26 |
| TAT | -6,05 | -3,47 | -6,1 | -2,7 | -4,36 | -5,23 | -0,15 | -3,28 | -4,8 | -4,23 | -4,02 | -2,38 | -0,67 | -2,64 | -0,57 | -2,34 | 0,25 | -1,84 | -3,88 | -4,01 | -0,46 | -2,98 | -2,08 | -5,61 | -2,36 | -2,79 | 1,34 | 0,36 | 0,89 | -2,41 | 0,14 | -4,79 | 1,99 | -1,88 | -2,37 | -6,4 | -0,96 | -2,18 | -4,14 | -2,96 | -1,78 | -1,61 | -0,37 | 0,4 | -0,31 | 0,84 | -1,39 | -2,03 | 0 | -3,76 | -2,2 | -0,74 | -4,57 | -2,76 | -4,85 | -0,38 | 0 | -2,09 | -3,89 | -1,39 | -4,22 |
| TAC | -2,97 | 2,16 | -1,28 | -2,16 | -2,6 | 1,13 | -1,04 | -0,37 | -0,56 | 1,04 | -0,05 | 2,76 | 4,31 | -0,66 | 1,17 | -0,67 | 3,1 | 1,53 | -1,68 | -3,74 | -3,38 | -1,71 | -1,59 | -1,1 | -0,25 | 0,47 | 2,89 | 2,5 | -1,17 | -2,32 | 1,04 | -5,24 | 1,42 | 0,15 | 1,04 | 2,05 | -0,77 | 0,84 | -4,08 | -0,03 | 0,83 | 3,34 | 5,51 | 2,24 | 0,56 | 1,28 | 1,18 | 0,14 | -3,05 | 0,06 | -4,5 | 2,15 | -0,66 | -3,34 | -3,3 | -1,98 | -1,5 | 2,98 | -0,63 | -1,98 | -3,03 |
| CAT | -7,07 | -2,89 | -2,89 | -6,32 | -4,53 | -5,4 | -3,4 | -3,47 | -4,71 | -0,75 | -2,07 | -2,23 | -0,7 | -0,67 | -0,36 | -2,44 | -1,58 | -1,03 | -2,34 | -3,93 | -2,13 | -1,33 | -3,74 | -3,8 | -2,76 | -3,12 | 1,3 | -2,01 | -2,88 | -3,71 | -0,87 | -5,3 | -4,5 | -5,36 | -5,77 | -5,6 | -4,6 | -0,91 | -7,07 | -5,63 | -3,17 | -1,73 | -4,62 | 0,23 | -3,86 | -1,22 | -2,34 | -2,18 | -2,64 | -1,81 | -3,19 | -3,32 | -3,43 | -2,12 | -4,27 | -1,9 | 0 | -0,66 | -1,38 | -3,88 | -4,07 |
| CAC | -0,14 | 2,89 | -2,51 | -1,53 | 0,63 | -1,27 | 0,83 | 1,65 | 0,26 | 4,28 | -1,9 | 2,92 | 2,82 | -2,74 | 2,61 | 1,85 | 2,01 | 0,71 | -0,71 | -2,98 | -2,18 | -1,18 | -3,78 | -2,47 | 0,71 | 1,83 | 3,03 | 2,71 | -1,38 | -2,66 | 2,02 | -4,06 | 0,73 | 1,62 | -3,08 | 0,6 | -2,17 | 2,22 | -1,45 | -1,44 | -2,49 | 0,76 | 2,78 | 0,29 | -3,71 | 0,29 | -0,07 | -3,07 | -0,99 | -0,8 | -5,09 | 1,51 | 0,31 | -1,88 | -4,2 | -2,67 | -0,39 | 1,32 | 0,33 | -0,08 | -3,21 |
| CAA | -6,4 | 0,22 | -5,78 | -4,08 | -6,92 | -4,62 | -3,86 | -4,74 | -5,69 | -3,08 | -2,94 | -4,06 | 2 | -0,35 | -1,51 | -2,62 | -1,69 | -1,88 | -1,98 | -4,8 | -6,12 | -5,26 | -1,29 | -3,49 | -3,04 | -5,81 | -3,14 | -4,59 | -4,55 | -7,25 | 1,06 | -2,49 | -3,68 | -2,07 | -3,68 | -1,97 | -8,14 | -2,26 | -5,27 | -5,25 | -4,86 | -4,03 | -4,01 | -2,37 | -0,26 | 0,35 | -0,4 | -0,02 | -3,02 | -4,35 | -3,76 | -2,95 | -6,92 | -5,22 | -7,2 | -1,82 | -1,85 | -1,49 | -3,4 | -3,92 | -3,68 |
| CAG | -5,87 | 0,66 | -5,09 | -5,27 | -7,09 | -2,88 | -6,18 | 2,53 | -1,59 | 0,35 | -5,53 | -1,29 | 2,07 | -3,44 | 0,31 | -3,16 | 0,88 | 3,12 | -4,73 | -4,1 | -1,41 | -1,65 | -2,61 | -3,23 | -3,23 | -2,8 | 5,76 | 1,66 | -2,25 | -5,36 | 1,4 | -5,07 | -3,32 | -3,3 | -2,46 | 0,76 | -6,28 | -2,48 | -8,19 | -3,66 | -2,98 | 2,14 | 4,09 | 0,19 | -3,06 | 0,17 | -0,18 | -3,58 | -3,45 | 0,37 | -2,89 | 0,37 | -2,32 | -6,85 | -8,23 | -2,83 | -1,63 | 1,28 | 0,55 | -3,78 | -5,94 |
| AAT | -2,59 | -4,46 | -6,15 | -5,21 | -3,71 | -4,76 | -2,11 | -4,98 | -6,19 | -2,41 | -0,26 | -1,05 | -1,94 | -3,48 | -0,31 | -2,72 | 0,38 | -2,18 | -2,56 | -1,17 | -2,76 | -3,91 | -2,71 | -4,73 | -1,63 | -2,73 | -2,05 | -1,5 | -2,37 | -5,28 | -0,26 | -4,74 | -1,01 | -4,47 | -3,8 | -2,79 | -4,46 | -4,11 | -3,92 | -6,34 | -5,58 | -2,2 | -4,58 | -3,67 | -2,46 | -0,17 | -2,07 | -1,19 | -1,38 | -4,76 | -1,27 | -2,5 | -4,39 | -1,42 | -3,1 | -1,22 | -2,33 | 0,18 | -2,02 | -1,67 | -5,17 |
| AAC | -2,22 | 5,88 | -3,19 | -0,54 | 0,32 | -1,18 | 2,07 | 2,95 | 1,2 | 5,88 | -2,37 | 4,01 | 2,94 | 1,22 | 4,08 | 3,76 | 5,6 | 5,31 | -2,18 | -2,23 | -3,72 | 0,6 | -2,01 | -4,64 | 0,76 | 4,56 | 1,55 | 2,67 | -2,74 | 0,22 | 2,13 | -0,31 | 4,17 | 0 | -0,03 | 1,03 | -1,93 | 3,38 | -0,99 | 0,16 | -1,73 | 4,08 | 5,49 | 3,91 | -0,55 | 2,6 | 4,3 | -0,7 | -1,25 | 0,24 | -3,76 | 3,55 | -2,57 | -3,95 | -4,41 | 0,6 | -1,02 | 3,79 | 0,59 | -2,23 | -3,73 |
| AAA | 1,04 | 4,9 | -3,22 | -3,72 | -3,08 | -0,9 | -1,86 | 7,12 | 1,56 | 4,38 | -0,86 | -0,17 | 7,48 | 1,18 | 4,56 | 3,2 | 6,35 | 2,8 | -1,97 | -2,29 | -4,38 | -2,51 | 0,25 | -4,47 | 2,01 | 0,66 | 3,01 | 2,58 | -3,45 | -3,1 | 7,77 | 0,48 | 8,37 | 3,9 | -0,97 | 2,54 | -2,44 | 2,77 | -0,13 | -0,81 | -1,57 | 1,97 | 4,25 | -1,61 | 3,84 | 7,18 | 7,31 | 1,8 | 0,62 | 1,43 | 1,53 | 4,12 | -1,56 | -4,25 | -5,16 | -5,11 | -3,51 | 6,6 | 5,68 | -2,05 | -3,42 |
| AAG | -3,48 | 1,82 | -1,99 | -1,81 | -0,36 | -2,28 | -2,69 | 2,26 | 0,52 | 3,45 | -2,66 | 0 | 2,33 | -0,39 | -2,53 | -2,26 | 4,56 | 0,85 | 0,5 | -0,25 | -1,98 | 4,98 | 0,34 | -4,62 | 0,06 | 0,4 | 3,45 | -0,35 | 0,23 | 0,65 | 2,31 | -1,15 | -0,62 | -1,8 | 1,8 | 2,9 | -0,95 | 2,67 | -2,33 | 0,17 | 1,77 | 1,32 | 6,24 | 3,31 | -1,87 | -0,21 | -0,22 | -2,41 | 0,61 | 0,67 | -1,1 | 2,5 | 0,41 | -3,69 | -3,36 | -0,17 | -1,21 | 0,39 | -0,2 | 0,03 | -1,84 |
| GAT | -5,9 | -0,42 | -5,9 | -5,05 | -1,23 | -4,3 | -2,23 | 2,69 | -4,54 | -0,84 | -3,22 | -0,04 | -0,56 | -2,35 | -1,37 | 0,25 | 0,78 | 0,79 | -1,19 | -2,82 | -3,34 | -1,76 | -2,27 | -5,36 | -0,14 | -0,03 | 2,01 | 0,24 | -1,85 | -3,84 | 1,79 | -2,51 | 1,84 | -1,03 | -4,67 | 0,33 | -2,46 | -0,85 | -1,82 | 0,05 | -4,77 | -0,94 | 1,75 | -2,07 | -2,77 | -0,58 | -2,45 | -3,35 | -1,08 | -1,93 | -0,43 | 0,9 | -3,37 | -1,96 | -3,67 | -1,51 | -1,19 | 1,3 | 0,93 | -3,49 | -1,65 |
| GAC | 1,12 | 4,45 | -2,78 | 1,41 | 1,55 | -0,01 | 0,82 | 5,04 | 0,25 | 6,3 | -0,37 | 3,01 | 6,11 | 1,44 | 5,51 | 2,51 | 1,33 | 1,35 | -0,99 | -2,64 | -3,41 | -1,7 | -2,54 | -2,15 | -1,57 | -0,57 | 2,38 | -0,07 | -2,2 | -1,62 | 2,89 | -1,22 | 3,26 | 0,64 | -0,76 | 1,3 | 2,19 | 3,05 | -0,98 | 2,46 | -1,2 | 2,48 | 5,4 | 2,32 | -2,54 | 1,74 | 4,02 | 1,76 | 0,64 | 1,98 | -4,1 | 6,59 | 0,8 | -0,78 | -1,56 | -3,42 | -2,88 | 7,04 | 3,29 | -3,19 | -3,97 |
| GAA | -0,93 | 6,61 | -3,61 | -4,57 | -3,93 | -2,13 | -2,68 | 4,83 | 0,12 | 4,72 | -1,44 | 2,67 | 7,23 | 0,21 | 3,64 | 0,71 | 4,8 | 1,82 | -3,87 | -3,41 | -5,97 | -3,31 | -0,03 | -3,83 | -0,22 | -1,44 | 1,79 | 2,45 | -2,81 | -3,51 | 5,01 | 0,19 | 4,12 | 3,12 | 0,24 | 4,38 | -3,58 | 2,01 | -4,23 | -2,98 | -2,55 | 1,7 | 4,54 | -5,46 | 0,9 | 5,12 | 5,45 | 2,26 | 1,87 | -3,75 | -4,65 | 4,18 | -3,14 | -1,73 | -5,47 | -1,55 | -4,04 | 7,26 | 5,69 | -3,91 | -3,36 |
| GAG | -3,62 | 3,93 | -4,89 | -4,11 | -0,44 | -1,8 | -4,59 | 2,56 | -5,65 | 2,66 | -2,08 | -1,54 | -4,15 | -3,02 | -1,37 | -4,33 | 2,4 | 1,13 | -1,05 | -3,14 | -3,38 | -2,26 | -0,99 | -3,66 | -2,76 | 1,15 | -0,43 | -1,91 | -2,56 | -2,52 | -3,92 | -3,34 | -3,49 | -6,7 | -2,31 | 0,19 | -4,05 | 1,35 | -3,24 | -1,9 | -4,41 | 2,82 | 1,21 | 2,24 | -3,11 | -0,96 | -4,64 | -3,82 | -2,66 | -1,26 | -2,53 | 0,82 | -0,7 | -2,75 | -4,83 | -2,31 | -2 | -0,79 | -3,43 | -4,91 | -3,53 |
| TGT | -4 | -1,36 | -1,49 | -2,33 | -2,26 | -2,89 | 0 | -3,65 | -3,23 | 0,96 | 0,26 | -0,22 | -0,64 | -3,21 | -1,72 | -0,95 | -0,34 | -0,8 | -0,39 | -1,79 | -3,56 | -3,66 | 0,29 | -2,15 | -1,4 | -1,59 | 3,3 | -2 | 1,6 | 0,63 | -0,71 | -0,97 | -1,52 | -0,14 | -4,39 | -1,95 | -2,05 | 1,07 | -2,96 | -3,05 | -1,12 | -1,74 | 0,59 | -0,59 | -2,24 | 0,42 | -0,69 | -1,82 | -0,15 | -0,12 | -3,48 | 0,1 | -2,17 | -0,41 | -2,75 | 0 | -1,97 | -0,36 | 0,42 | -3,48 | -2,16 |
| TGC | -3,38 | 0,72 | -1,8 | -1,7 | -2,58 | 0,25 | 1,28 | -1,81 | -0,55 | 0,66 | -2,79 | -2,14 | -0,75 | -2,57 | 1,62 | -2,59 | 0,88 | 2,07 | -0,35 | -1,04 | -0,41 | -1,05 | -3,23 | -1,23 | 0,63 | 0,84 | -0,4 | 1,1 | -0,61 | -1,46 | -1,27 | -4,98 | 0,39 | -2,85 | -3,17 | -0,64 | -3,87 | -0,21 | -3,8 | -3,25 | -2,86 | -1,48 | 0,12 | 1,3 | -2,32 | 1 | 1,57 | -0,78 | -1,77 | -1,85 | -3,02 | 1,55 | 0,17 | -0,12 | -3,42 | 0,82 | -1,39 | -0,72 | -1,71 | -2,85 | -5,31 |
| TGG | -8,04 | 1,6 | -5,34 | -7,67 | -5,63 | -3,47 | -8,01 | -9,17 | -5,51 | -1,3 | -4,37 | -3,78 | 0,83 | -2,46 | -3,08 | -3,59 | 0,93 | 0,6 | -3,19 | -5,69 | -5,77 | -1,51 | -3,93 | -3,9 | -3,93 | -4,81 | 0,12 | -0,8 | -2,68 | -1,81 | -1,83 | -3,64 | -3,1 | -4,78 | -4,96 | -0,81 | -4,29 | -2 | -9,04 | -7,69 | -3,99 | 0,3 | 0,07 | -2,5 | -2,45 | -1,18 | -2,05 | -3,68 | -3,08 | -4,1 | -5,45 | -4,44 | -5,98 | -7,23 | -7,51 | -0,59 | -1 | -0,18 | -3,1 | -4,95 | -3,79 |
| CGT | -4,9 | 2,92 | -5,67 | -2,96 | -4,02 | -0,24 | -1,81 | 2,35 | -0,13 | 6,02 | -1,45 | 1,47 | 4,49 | -0,17 | 4,17 | -0,89 | 4,71 | 2,98 | -1,97 | -5 | -2,4 | -0,33 | 4,7 | -3,07 | -0,69 | 1,76 | 6,22 | 5,23 | -1,6 | -2,73 | 7,5 | -1,04 | 2,6 | 1,04 | -3,42 | 0,8 | -1,77 | 2,7 | 0,12 | 1,99 | -2,25 | 6,42 | 5,01 | 3,93 | 1,27 | 3,42 | 3,92 | 1,89 | -3,3 | -2,36 | -3,91 | 7,11 | -0,69 | -1,8 | -6,04 | 0,11 | -1,87 | 4,35 | 1,41 | -3,05 | -2,14 |
| CGC | -5,9 | 1,49 | -6,37 | -4,82 | -2,88 | -0,57 | -1,16 | -2,29 | -1,81 | -1,46 | -2,62 | -1,83 | 4,6 | -3,12 | 4,14 | -3,23 | -0,47 | 0,76 | -3,34 | -3,14 | -5,79 | -5 | -2,24 | -0,07 | -3,88 | -2,21 | -1,55 | -3,01 | -5,56 | -4,51 | 3,16 | -4,48 | -0,16 | -3,14 | -4,66 | -1,92 | -5,19 | -3,06 | -2,53 | -6,28 | -4,67 | -0,55 | -2,89 | 1,16 | -2,37 | -0,75 | -2,04 | -3,77 | -4,52 | -0,98 | -7,37 | -2,28 | -2,8 | -2,92 | -3,1 | -3,06 | -4,06 | 3,12 | -2,99 | -2,71 | -5,78 |
| CGA | -2,23 | -1,47 | -5,73 | -5,38 | -2,93 | -5,04 | -1,22 | -7,53 | -3,16 | -1,55 | -2,83 | -5,2 | -1,69 | -2,08 | 0,19 | -2,7 | 0,1 | -0,05 | -0,65 | -2,75 | -2,51 | -2,61 | -3,88 | -3,38 | -3,57 | -3,54 | -1,34 | -3,34 | -3,4 | -5,09 | -1,39 | -0,73 | -1,37 | -3,92 | -2,76 | -2,49 | -4,13 | -3,85 | -3,54 | -6,14 | -5,6 | -2,53 | -5,94 | -2,62 | -0,2 | -0,86 | -5,41 | -3,19 | 0,52 | -1,86 | -1,89 | -3,89 | -4,97 | -1,72 | -2,79 | -2,76 | 0 | -0,44 | -2,88 | 0 | -2,28 |
| CGG | -4,56 | -0,19 | -4,38 | -6,1 | -5,07 | -2,41 | -2,57 | -10,85 | -3,69 | -3,11 | -1,5 | -6,12 | -5,24 | -2,67 | -5,21 | -7,03 | -1,34 | 0 | -1,72 | -0,29 | -0,27 | -1,03 | -1,32 | -2,48 | -1 | -2,33 | -2,14 | -2,2 | -2,54 | -4,7 | -2,43 | -3,91 | -5,93 | -8,51 | -2,98 | -2,77 | -6 | -2,85 | -7,3 | -5,84 | -0,67 | -0,17 | -4,28 | 1,86 | -3,51 | -2,77 | -6,23 | -3,96 | -2,48 | -1,41 | -1,86 | -5,49 | -7,22 | -3,83 | -4,82 | 0 | -2,49 | -1,82 | -2,05 | -1,1 | -3,03 |
| AGA | -2,01 | -0,77 | -2,1 | -1,85 | -0,45 | 0,86 | -2,24 | -4 | -0,79 | -1,82 | -1,41 | -3,27 | 0,31 | -3,5 | -1 | -1,05 | 0 | 1,35 | -1,45 | -1,41 | -0,33 | -0,9 | 0,29 | -0,79 | -2,3 | -2,11 | 2,49 | -2,69 | -2,57 | -1,46 | -3,27 | 0,83 | -4,04 | -2,25 | 1,69 | -1,87 | -2,04 | -0,97 | -2,17 | -2,7 | -0,92 | -1,97 | -0,82 | -3,1 | -0,33 | -0,79 | -1,18 | -0,83 | 0 | -1,9 | -2,75 | -1,91 | -2,56 | 0,67 | -1,8 | -1,75 | -1,4 | -1,25 | -1,06 | -0,77 | -1,68 |
| AGG | -2,68 | -0,43 | -1,5 | -0,56 | -1,94 | -2,55 | -2,8 | -4,71 | -1,62 | -1,26 | -0,16 | -1,99 | -1,79 | -2,66 | -1,3 | -2,18 | 0,5 | 0 | -0,28 | 0,26 | -1,76 | -2,09 | 0,58 | -2,15 | -1,2 | -2,69 | -0,16 | -1,69 | -1,79 | -2,7 | -0,29 | -1,91 | -1,88 | -3,36 | -1,52 | -1,68 | -1,95 | -2,1 | -2,72 | -2,39 | 0,08 | -0,7 | -4,01 | 0,01 | -2,56 | -2,79 | -0,94 | -0,09 | 0 | 0 | -2,17 | -0,3 | -1,86 | -2,64 | -1,97 | -2 | -2,05 | -0,82 | -1,44 | -2,94 | -0,54 |
| GGT | -1,22 | 5,75 | -2,56 | -2,91 | -0,9 | -1,88 | -0,24 | 4,21 | -0,36 | 4,96 | -2,64 | 3,58 | 4,2 | -1,76 | 2,57 | -0,79 | 8,87 | 1,23 | -0,66 | -2,35 | -1,73 | 1,1 | 0,8 | 0,67 | 0,31 | 3,86 | 9,8 | 3,14 | -0,39 | 0,67 | 3,81 | -0,36 | 2,37 | -0,58 | -3,43 | 1,45 | -0,6 | 1,74 | -1,99 | 3,98 | -1,95 | 5,88 | 7,26 | 3,98 | 0,98 | 4,29 | 2,96 | -2,51 | -0,17 | 1,75 | -2,33 | 9,4 | 1,52 | -1,79 | -3,06 | -1,39 | 0,27 | 3,98 | 1,86 | -1,86 | -2,05 |
| GGC | -4,42 | 2,72 | -6,66 | -3,91 | -1,35 | -0,48 | -4,05 | 1,76 | -4,03 | 3,14 | -4,36 | 0,12 | 1,28 | -4,09 | 1,62 | -2,09 | -0,05 | -0,18 | -1,3 | -3,13 | -5,87 | -2,59 | 0,26 | -3,46 | 1,43 | 1,37 | 2,55 | 0 | -1,89 | -2,79 | 4,34 | 1,67 | 0,3 | -2,03 | -5,26 | 0,28 | 1,55 | 2,68 | -0,46 | 2,96 | -3,86 | 1,36 | 1,96 | -1,35 | -1,35 | 2,68 | 4,47 | -3,54 | 1,08 | 0,03 | -6,65 | 6,64 | 2,49 | -1,07 | 0,01 | -4,11 | -2,19 | 1,76 | -0,23 | -4,14 | -5,92 |
| GGA | -6,46 | -1,87 | -7,03 | -5,7 | -2,86 | -1,02 | -1,37 | -2,35 | -5,14 | -1,19 | -2,63 | -5,19 | -2,65 | -0,32 | -1,74 | -4,86 | -0,85 | -0,97 | -0,81 | -3,06 | -3,15 | -0,97 | -2,49 | -2,28 | -0,59 | -4,76 | -0,38 | -2,3 | -3,56 | -4,26 | -0,99 | -1,49 | -3,98 | -3,76 | -2,45 | 0,24 | -2,41 | -0,83 | -3,67 | -1,63 | -4,11 | -2,87 | -3,96 | -1,78 | -3,04 | -1,62 | -5,25 | -4,75 | -2,73 | 0,11 | -3,12 | -3,41 | -2,99 | -2,27 | -3,76 | -2 | -3,28 | -1,44 | -3,84 | -2,77 | -0,44 |
| GGG | -3,32 | -1,19 | -5,16 | -6,48 | -5,31 | -0,22 | -5,74 | -9,01 | -7,42 | -3,26 | -4,07 | -6,89 | -1,36 | -2,84 | -4,32 | -1,98 | 2,92 | -2,41 | -1,07 | -1,52 | -0,99 | -1,09 | -1,91 | -2,02 | -2,49 | -4,31 | 1,2 | -0,25 | -3,93 | -3,5 | 0,27 | -1,45 | -1,64 | -6,86 | -4,47 | -1 | -3,92 | -2,45 | -6,63 | -6,81 | -2,68 | 0,31 | -4,33 | -0,74 | -2,14 | -0,69 | -4,55 | -2,68 | 0,38 | -1,6 | 0,6 | -4,47 | -6,22 | -5,36 | -6,61 | 0,32 | -1,16 | -0,54 | -2,07 | -1,86 | -1,36 |
